# Neuronal polyunsaturated fatty acids are protective in FTD/ALS

**DOI:** 10.1101/2024.01.16.575677

**Authors:** A Giblin, AJ Cammack, N Blomberg, A Mikheenko, M Carcolé, R Coneys, L Zhou, Y Mohammed, D Olivier-Jimenez, ML Atilano, T Niccoli, AN Coyne, R van der Kant, T Lashley, M Giera, L Partridge, AM Isaacs

## Abstract

We report a conserved transcriptomic signature of reduced fatty acid and lipid metabolism gene expression in human post-mortem ALS spinal cord and a *Drosophila* model of the most common genetic cause of FTD/ALS, a repeat expansion in *C9orf72*. To investigate lipid alterations, we performed lipidomics on C9FTD/ALS iPSC-neurons and post-mortem FTLD brain tissue. This revealed a common and specific reduction in phospholipid species containing polyunsaturated fatty acids (PUFAs). To determine whether this PUFA deficit contributes to neurodegeneration, we fed C9FTD/ALS flies PUFAs, which yielded a modest increase in survival. However, increasing PUFA levels specifically in neurons of the *C9orf72* flies, by overexpressing fatty acid desaturase enzymes, led to a substantial extension of lifespan. Neuronal overexpression of fatty acid desaturases also suppressed stressor induced neuronal death in C9FTD/ALS patient iPSC-neurons. These data implicate neuronal fatty acid saturation in the pathogenesis of FTD/ALS and suggest that interventions to increase PUFA levels specifically within neurons will be beneficial.

## Introduction

Amyotrophic lateral sclerosis (ALS) and frontotemporal dementia (FTD) are two relentlessly progressive and invariably fatal neurodegenerative disorders. ALS is characterized by loss of upper and lower motor neurons in the brain and spinal cord, leading to muscle wasting and paralysis, while FTD leads to degeneration of the frontal and temporal lobes of the brain, resulting in behavioural and language abnormalities. It is now well established that ALS and FTD represent two ends of a disease continuum, with overlapping clinical and pathological features. ALS and FTD are also linked genetically, with the most common genetic cause of both diseases being an intronic G_4_C_2_ repeat expansion in the *C9orf72* gene (C9FTD/ALS) (*1, 2*).

The *C9orf72* repeat is transcribed bidirectionally into sense and antisense repeat RNAs, which are translated into dipeptide repeat proteins (DPRs) by a process termed repeat-associated non-ATG (RAN) translation (*3–8*). RAN translation occurs in all reading frames and on both strands to produce five distinct DPR species: poly(GR), poly(GP) and poly(GA) from the sense strand, and poly(GP), poly(PR) and poly(PA) from the antisense strand. DPRs and the repetitive RNAs themselves have been implicated in driving neurodegeneration (*9–15*). In addition, the repeat expansion leads to reduced levels of the C9orf72 protein (*16, 17*). Functional and genetic evidence suggests that loss of function is insufficient to cause neurodegeneration on its own (*18–20*), but plays a role through exacerbating gain of function mechanisms (*21, 22*). Despite numerous cellular pathways implicated downstream of the *C9orf72* repeat expansion since its discovery (*23, 24*), the molecular mechanisms driving neuronal loss are still unclear.

The brain has the second highest lipid content of any organ in the body, with these molecules serving as critical components of neuronal and organellar membranes. Brain lipids contain a particularly high proportion of polyunsaturated fatty acids (PUFAs) (*25*), and epidemiological studies have demonstrated that increased dietary consumption of PUFAs, particularly omega-3 PUFAs, are associated with decreased ALS risk and longer survival after onset (*26–28*). However, a molecular understanding of these findings and their relevance to neurodegeneration are unclear. Thus, in this study we sought to characterize lipid changes associated with C9FTD/ALS and understand their contribution to neurodegeneration.

## Results

### Downregulation of fatty acid and lipid metabolism pathways in C9FTD/ALS

To identify pathways dysregulated in neurons in response to expression of the pathologic *C9orf72* repeat (C9) expansion, we performed RNA sequencing (RNA-seq) on *Drosophila* heads with 36 G_4_C_2_ repeats expressed exclusively in adult neurons (*9*). These experiments were performed at an early timepoint (5 days of repeat expression) to assess early gene expression changes. We found widespread dysregulation of lipid metabolism genes in C9 flies, including multiple genes in the canonical fatty acid synthesis and desaturation pathway (**Fig. 1A**) such as *AcCoAS, FASN1, FASN2* and *Desat1* (**Fig. 1B, Sup Fig. 1A**). Gene ontology (GO) enrichment analysis of differentially expressed genes identified only three GO terms enriched among upregulated pathways **(Sup Fig. 1B**). However, multiple terms related to fatty acid and lipid metabolism were among the most significantly downregulated pathways (**Fig. 1C**). To determine whether these lipid gene expression changes were conserved in human disease, we re-analyzed the largest bulk RNA-seq dataset generated from ALS post-mortem spinal cords, comprising 138 ALS cases and 36 non-neurological disease controls (*29*). Strikingly, genes in the same lipid and fatty acid metabolism pathway were also downregulated in ALS spinal cords, including *ACACA*, *ACSS2*, *FASN*, *ELOVL6* and *SCD* (orthologous to *Drosophila ACC, AcCoAS*, *FASN1*/*FASN2*, *Baldspot* and *Desat1* respectively) (**Fig. 1D**). These genes were similarly downregulated in the 28 C9 ALS patient spinal cords present in the dataset **(Sup Fig. 2**), suggesting this pathway is broadly dysregulated in ALS. Together, these findings demonstrate conserved transcriptional dysregulation of lipid metabolism, and specifically downregulation of fatty acid synthesis and desaturation processes, in FTD/ALS neurons.

**Figure 1.**
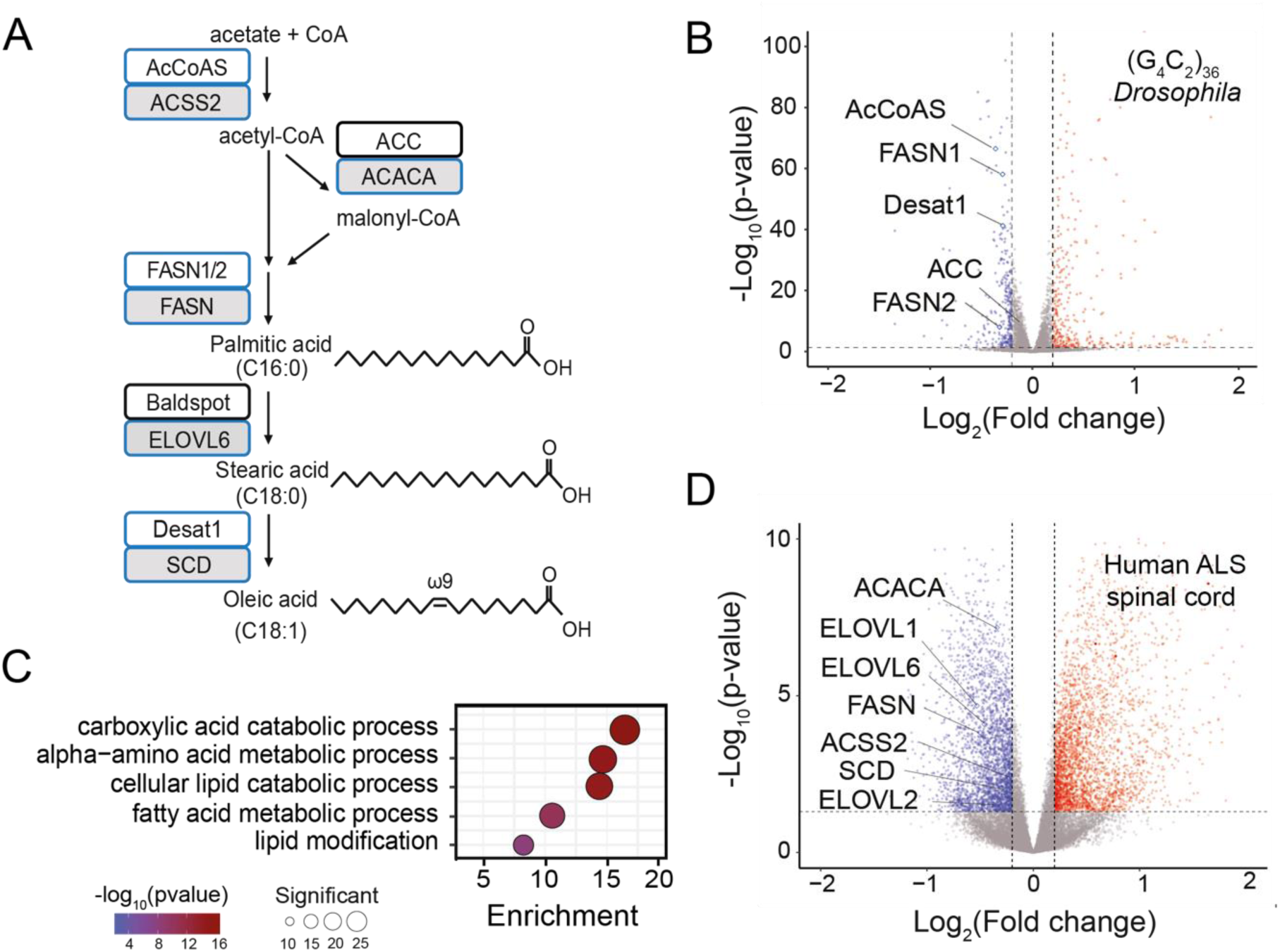
Transcriptomic analysis reveals downregulation of fatty acid and lipid metabolism genes in C9ALS/FTD. (A) Simplified lipid synthesis and desaturation pathway, with *Drosophila* genes in white boxes and human orthologous genes in grey boxes. Blue boxes indicate genes that were significantly downregulated in (B) and (D). The “C” number indicates the number of carbons in the fatty acyl chain, the number after the colon denotes the number of double bonds, and the “ω” number denotes the position of the final double bond before the methyl carbon. (B) Volcano plot highlights significantly downregulated fatty acid synthesis and desaturation genes in C9 flies. (C) Gene ontology (GO) biological process enrichment analyses showing lipid metabolism terms significantly enriched among downregulated genes in C9 flies. (D) Volcano plot of RNA-seq data from cervical spinal cord of 138 individuals with ALS (including C9 ALS) and 36 non-neurological disease controls from the New York Genome Center ALS Consortium (*29*), highlighting downregulated canonical fatty acid synthesis and desaturation pathway genes.

**Figure 2.**
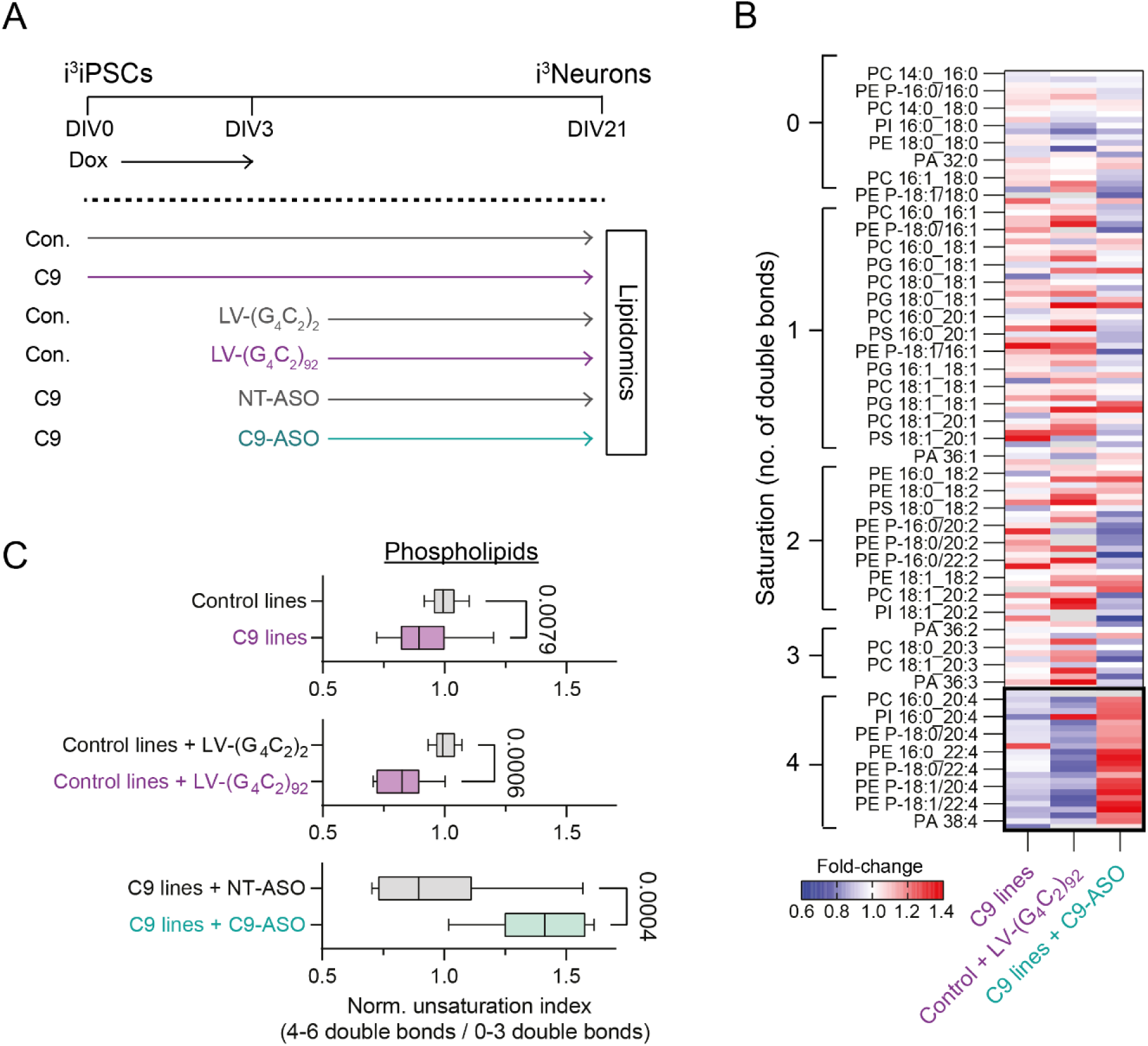
C9 repeats cause loss of highly unsaturated phospholipid species in iPSC-derived neurons. (A) C9 and isogenic control i^3^iPSCs were induced with the i^3^Neuron protocol (*30, 31*) and cultured for 21 days in vitro (DIV21) for lipidomic analyses. To confirm disease-specificity of lipid changes, control lines were transduced with (G_4_C_2_)_92_ repeat lentivirus (or (G_4_C_2_)_2_ control), and C9 lines were treated with a *C9orf72* antisense oligonucleotide (ASO) or a non-targeting (NT) control (*7*). (B) Heatmap displaying all detected phospholipids as average fold-change over control across C9 lines and conditions (n=3 C9 lines, n=9 neuronal inductions; n=2 control lines+lentiviruses, n=3 neuronal inductions; n=3 C9 lines+ASO, n=6 neuronal inductions). Highly unsaturated species (≥4 double bonds, outlined) were reduced in C9 and LV-(G_4_C_2_)_92_ lines but increased in C9 lines with the C9-ASO, indicating that these changes were specific to C9 repeats. Lipids were normalized by lipid class (see Methods for details on filtering and normalization). Grey boxes indicate phospholipid species which were outside the fold-change range. (C) Quantification of normalized unsaturation index across C9 and control lines and conditions. Each index is normalized to its own control within each neuronal induction; thus, the average of each control condition is 1. The index was decreased in C9 lines compared to control lines and in control lines expressing 92 repeats compared to 2-repeat control, but increased in C9 lines when administered the C9-ASO compared to NT-ASO control. *Top:* n=17 across 2 isogenic control lines; n=24 across 3 C9 lines; *Middle:* n=8 per lentivirus across 2 control lines; *Bottom:* n=15 for each ASO treatment across 3 C9 lines. Lines and error bars represent median and minimum/maximum, respectively, and significance was calculated with two-tailed, unpaired Student’s t-test.

### Phospholipid saturation is dysregulated in *C9orf72* iPSC-derived neurons

In light of the dysregulation of fatty acid and lipid metabolism transcriptional pathways, we next determined whether lipids were altered in C9 neurons. We performed lipidomic analyses on C9 repeat-containing iPSC-cortical neurons and isogenic controls, which were induced with the i^3^Neuron cortical protocol (*30, 31*), and collected 21 days later (**Fig. 2A**). Among the different lipid classes that were measured, these experiments revealed a consistent change across phospholipid species. Phospholipids are composed of two fatty acyl chains and a head group. The number of double bonds in each chain determines their saturation with 0 double bonds being completely saturated and each additional double bond increasing unsaturation, with ≥2 double bonds classed as polyunsaturated. We observed a striking shift towards higher phospholipid saturation and loss of highly polyunsaturated phospholipids (containing fatty acyl chains with ≥4 double bonds) compared to controls (**Fig. 2B-C; Sup Fig. 3 and 4A-C)**. To confirm that these changes were driven by the C9 repeat expansion and not cell line variability or other mechanisms such as *C9orf72* loss of function, we next performed lipidomic analyses in two cross-validation experiments (**Fig 2A**). In the first, we exogenously expressed C9 repeats in control iPSC-neurons by transducing with (G_4_C_2_)_92_ or (G_4_C_2_)_2_ lentiviruses. As expected, lentiviral repeat expression resulted in substantial DPR production in (G_4_C_2_)_92_ but not (G_4_C_2_)_2_-transduced neurons **(Sup Fig. 5**). Exogenous repeat expression in control iPSC-neurons recapitulated the loss of highly polyunsaturated phospholipids we observed in the C9 patient lines (**Fig. 2B-C**). This suggests that expression of expanded *C9orf72* repeats is sufficient to drive the lipid changes observed. Next, in three C9 repeat lines we knocked down with an antisense oligonucleotide (ASO) that specifically targets transcripts containing the C9 repeat (*7*). This led to a near complete (>95%) reduction in DPRs compared to a non-targeting control, confirming effective knockdown **(Sup Fig. 5**). C9 repeat knockdown prevented the reduction in highly polyunsaturated phospholipids we observed in the C9 patient lines, suggesting that the C9 repeat was driving these changes (**Fig. 2B-C; Sup Fig. 4D-G**). To quantify these changes in saturation, we calculated a desaturation index as the ratio of highly unsaturated (4-6 double bonds) to less unsaturated species (0-3 double bonds) and observed a significantly reduced unsaturation index in the C9 lines and control lines expressing (G_4_C_2_)_92_, which was prevented by ASO treatment, thus confirming that phospholipid saturation is shifted by C9 repeats (**Fig. 2C**). Lipid class proportionality was similar across conditions, suggesting that these observations are due to a specific shift in phospholipid saturation rather than a global alteration in lipid class abundance **(Sup Fig. 6**). Together, these results demonstrate a striking and specific decrease in highly polyunsaturated phospholipid species caused by the presence of expanded C9 repeats.

**Figure 3.**
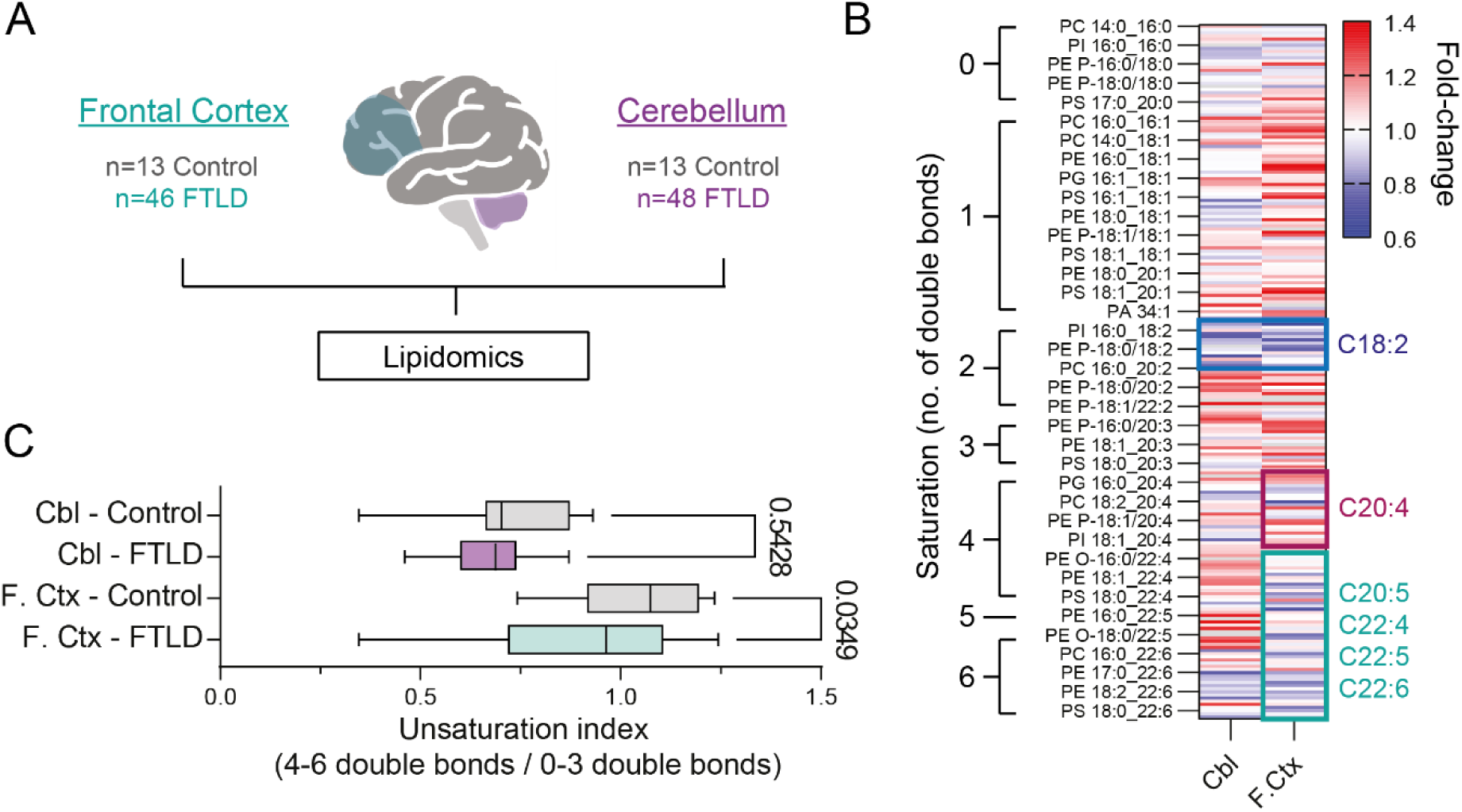
Highly unsaturated phospholipids are decreased in FTLD post-mortem frontal cortex. (A) Targeted lipidomics was done on post-mortem tissues from FTLD and age- and sex-matched control frontal cortex and cerebellum samples. (B) Heatmap displaying all detected phospholipids as fold-change over control in each brain region individually. Highly unsaturated species (≥4 double bonds, except C20:4) are outlined in cyan and show broad downregulation in FTLD frontal cortex but not cerebellum. Species containing arachidonic acid (C20:4) are outlined in red, and many display upregulation in FTLD frontal cortex. Species containing linoleic acid (C18:2) are outlined in blue and show downregulation in both tissue regions. (C) Unsaturation indices of phospholipids from FTLD and control brain regions, demonstrating a significant reduction in frontal cortex but not in cerebellum (One-way ANOVA F=23.24, p<0.0001; post-hoc comparisons of FTLD vs. control p-values adjusted with Šídák’s multiple comparisons test).

**Figure 4.**
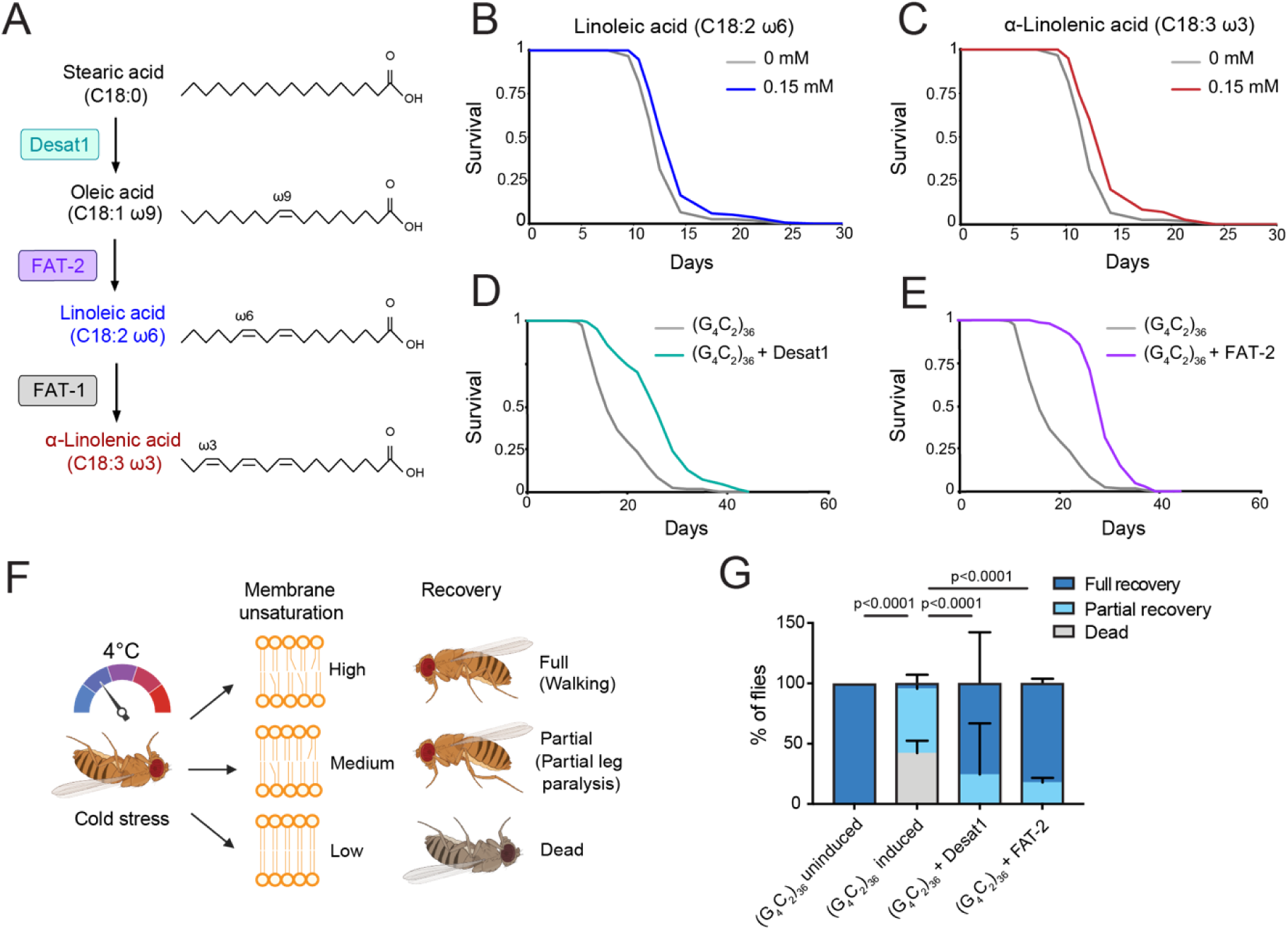
Promoting fatty acid desaturation through either genetic or feeding paradigms extends C9 fly survival and prevents cold stress-induced death and paralysis, an *in vivo* measure of membrane fluidity. (A) Simplified fatty acid desaturation pathway. (B-C) Dietary supplementation of linoleic or α-linolenic acid at 0.15mM extended C9 fly survival (Linoleic acid p=5.697×10^-5^; α-linolenic acid p=2.951×10^-6^). (D-E) Neuronal overexpression of *Desat1* or *FAT-2* extended C9 fly survival (*Desat1* p=5.839×10^-20^; *FAT-2* p=5.119x10^-38^). n=150 flies per condition. Log-rank test was used for all comparisons. (F) Schematic diagram of cold stress assay. Flies were exposed to 4°C, which induces a cold-induced paralysis response. Membrane unsaturation influences tolerance to cold stress, with higher membrane unsaturation providing greater tolerance. After 18 hours, flies were moved to room temperature for 1 hour, and recovery was assessed. (G) (G_4_C_2_)_36_ flies were sensitive to cold stress, with significantly increased death and paralysis, and decreased recovery compared to uninduced controls (p<0.0001). Neuronal overexpression of *Desat1* or *FAT-2* in (G_4_C_2_)_36_ flies significantly increased the proportion of flies experiencing a full recovery compared to (G_4_C_2_)_36_ alone and significantly reduced death post-exposure compared to (G_4_C_2_)_36_ alone. n=3 biological replicates, containing 15 flies per replicate vial. Results were analyzed by χ2 test. Data presented as mean ± S.D. Genotypes: UAS-(G_4_C_2_)_36_, elavGS; UAS-(G_4_C_2_)_36_, elavGS/UAS-Desat1; UAS-(G_4_C_2_)_36_, elavGS/UAS-FAT-2.

**Figure 5.**
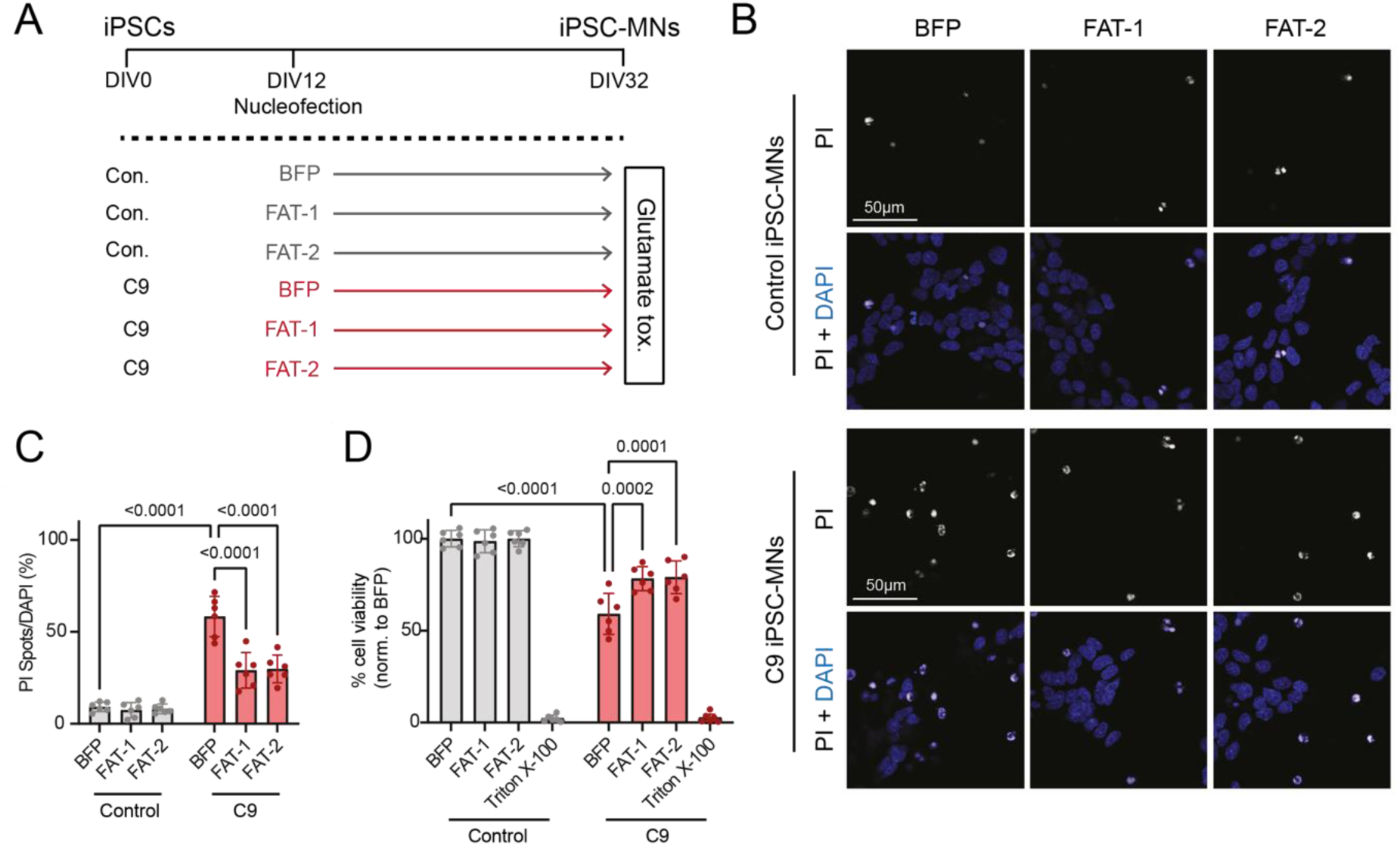
FAT-1 and FAT-2 rescue glutamate-induced excitotoxicity in C9 iPSC-derived spinal neurons. (A) Schematic of spinal neuron differentiation timeline, with timepoints of nucleofection and glutamate-induced excitotoxicity measurements. (B) Representative confocal images of cell death in control and *C9orf72* iPS-SNs expressing BFP, FAT1 or FAT2, as measured by propidium iodide (PI) incorporation. (C) Quantification of the ratio of propidium iodide (PI) positive spots to DAPI positive nuclei (quantification of cell death) following 4-hour exposure to 10 µM glutamate. n=6 control and n=6 C9 iPSC lines. Data points represent average percent cell death across 10 images per well. (D) Percent cell viability as measured by Alamar Blue following 4-hour exposure to 10 µM glutamate. n=6 control and n=6 C9 iPSC lines. Data points represent average percent viability from 3 replicate wells for each condition. Two-way ANOVA with Tukey’s multiple comparison test was used to calculate statistical significance. Data presented as mean ± S.D.

### Phospholipid saturation is dysregulated in FTLD post-mortem frontal cortex

We next asked whether phospholipid saturation dysregulation is also present in human disease tissue. We performed lipidomic analyses on post-mortem affected (frontal cortex) and less affected (cerebellum) brain tissue from a large cohort of 47 individuals with neuropathologically confirmed FTD, termed frontotemporal lobar degeneration (FTLD), 15 of whom had a C9 mutation, and 13 age- and sex-matched, healthy controls (**Fig. 3A**). In clear concordance with our iPSC-neuron data, in FTLD frontal cortex we observed a decrease in highly unsaturated phospholipids, particularly those containing ≥4 double bonds in their most unsaturated fatty acyl chain (**Fig. 3B-C; Sup Fig. 7**), while lipid class proportionality was similar between control and FTLD tissue in both brain regions **(Sup Fig. 8**). Interestingly, there was one exception, in species containing arachidonic acid (C20:4), some of which were upregulated in FTLD tissues. This is consistent with the association of arachidonic acid with inflammatory signalling (*32*), highlighting additional lipid alterations that occur in end-stage disease, as well as a previous study showing elevated arachidonic acid in C9 repeat disease models (*33*). These changes in highly polyunsaturated phospholipids were largely specific to the frontal cortex. Interestingly, in both tissue regions, we also observed a decrease in species containing linoleic acid (C18:2), an essential PUFA that is a precursor for highly unsaturated fatty acid species (*34*). Thus, consistent with C9 iPSC-neurons, FTLD post-mortem brains displayed a reduction in highly polyunsaturated species, specifically in the affected region.

### Promoting neuronal fatty acid desaturation increases the survival of C9 flies

To determine whether dysregulated lipid metabolism directly contributes to neurotoxicity, or is simply a bystander, we first asked whether dietary supplementation with fatty acids could rescue survival of C9 flies. The PUFAs linoleic acid (C18:2) and α-linolenic acid (C18:3) significantly but modestly extended median survival of C9 flies by 12-15% (**Fig. 4B-C, Sup Fig. 9A-C),** whereas adding saturated or monounsaturated fatty acid species (palmitic acid (C16:0), stearic acid (C18:0), and oleic acid (C18:1)) had either no effect or decreased survival **(Sup Fig. 9D-F**). The extensions in survival generated by PUFA supplementation were specific to the disease model, because supplementing wildtype flies with linoleic acid or α-linolenic acid either decreased or had no effect on wildtype survival **(Sup Fig. 10A-B**). Further, these rescues were not due to an alteration in feeding behaviour, as measured by the proboscis extension assay **(Sup Fig. 10C-D**).

As the rescue with feeding PUFAs was modest, we next asked whether neuronal overexpression of fatty acid synthase or desaturase genes, which encode enzymes that produce and desaturate long chain fatty acids respectively (**Fig. 4A**), could prevent C9-associated neurodegeneration *in vivo*. We crossed our C9 flies to flies overexpressing lipid pathway genes, using the same adult neuronal driver as the C9 repeats, and measured survival. Whilst overexpression of the fatty acid synthase genes *FASN1* and *FASN2* resulted in survival extensions **(Sup Fig. 11A-B**), the most impressive rescues occurred when overexpressing fatty acid desaturases. Overexpression of *Desat1*, which introduces a double bond into the acyl chain of saturated fatty acids to produce the monounsaturated fatty acid oleic acid, significantly extended C9 fly survival, increasing median survival from 15 days to 25 days, an increase of 67% (**Fig. 4D).** Linoleic acid (C18:2) and α-linolenic acid (C18:3) are termed essential fatty acids, because these species cannot be synthesized endogenously by most animals, including *Drosophila* and humans, and must be obtained from the diet to serve as precursors for generating highly polyunsaturated fatty acids. Interestingly, the nematode *Caenorhabditis elegans* does possess fatty acid desaturase genes, which encode enzymes capable of endogenously synthesizing these essential PUFAs. Neuronal expression in C9 *Drosophila* of *C. elegans FAT-2*, a Δ12/Δ15 fatty acid desaturase which produces linoleic acid and α-linolenic acid from monounsaturated fatty acids (*35, 36*), provided an even greater genetic rescue, extending median survival from 15 days to 27.5 days, an increase of 83% (**Fig. 4E**). Importantly, these genetic rescues were not due to an effect on DPR levels, because poly(GP) levels in C9 fly heads were unchanged by overexpression of any lipid-related genes **(Sup Fig. 11C-D**). Further, these survival extensions were largely specific to C9 disease, as overexpressing fatty acid synthases or desaturases in neurons of healthy control flies had either no effect on survival or increased survival by a much smaller magnitude than that observed in the context of C9 repeats **(Sup Fig. 12**). In summary, these results demonstrate that promoting lipid desaturation within neurons is particularly beneficial for ameliorating C9-associated neurodegeneration *in vivo*.

Lipid saturation influences the biophysical properties of cellular membranes, particularly the packing of membrane phospholipids, with increased desaturation resulting in an increase in membrane fluidity (*35, 37, 38*). Since membrane fluidity increases with temperature, poikilothermic *Drosophila* must adjust their membrane lipid content upon temperature fluctuations to survive (*39, 40*). Indeed, *Drosophila* alter their feeding preferences in response to cold exposure to incorporate more PUFAs into their lipid bilayers to maintain their membrane fluidity (*40*). Therefore, to investigate the mechanism by which neuronal desaturase expression is beneficial, and whether membrane fluidity plays a role, we used the *Drosophila* cold stress membrane fluidity paradigm. C9 flies were exposed to 4°C for 18 hours, which causes a cold-induced paralysis attributable to decreased membrane fluidity (*35, 41, 42*), and then returned to room temperature, with recovery scored one hour later (**Fig. 4F**). Whereas all non-repeat expressing control flies showed a full recovery after this period, 42% of flies expressing (G_4_C_2_)_36_ were dead, and 54% were partially paralysed, with only 2% exhibiting a full recovery (**Fig. 4G**). Strikingly, overexpressing either *Desat1* or *FAT-2* specifically in neurons prevented death entirely after cold exposure in C9 flies and resulted in a dramatically improved recovery (**Fig. 4G**). These data suggest an increase in membrane fluidity contributes to the beneficial effect of neuronal desaturase overexpression.

### *FAT-1* or *FAT-2* overexpression rescues stressor induced toxicity in C9 iPSC-derived spinal neurons

Finally, we asked whether overexpression of desaturase genes could prevent C9-associated neurodegeneration in human C9 neurons. We overexpressed the *C. elegans* lipid desaturases *FAT-1* or *FAT-2*, or a *BFP*-only control, in C9 iPSC-derived spinal neurons and then exposed them to high levels of glutamate to induce excitotoxicity (**Fig. 5A**). As previously reported (*10, 43*), C9 spinal neurons exhibited heightened susceptibility to excitotoxic cell death compared to spinal neurons derived from healthy donor iPSCs (**Fig. 5B-D**). Importantly, however, overexpression of either *FAT-1* or *FAT-2* was sufficient to partially rescue glutamate induced toxicity, significantly decreasing cell death in C9 spinal neurons compared to *BFP*-only control (**Fig. 5B-D**). Thus, desaturase overexpression is beneficial in preventing C9-associated neurodegenerative phenotypes in human C9 neurons.

## Discussion

In this study, we uncovered lipid metabolism dysregulation in multiple models of C9FTD/ALS, including transgenic *Drosophila,* iPSC-derived neurons and ALS and FTD patient post-mortem brain and spinal cord. We first identified transcriptional dysregulation of canonical fatty acid synthesis and desaturation genes, which was present at a pre-degeneration timepoint in C9 fly heads and was conserved in end stage disease in ALS. Through lipidomic assays in C9 iPSC-derived neurons and FTLD post-mortem brains, we identified a loss of highly polyunsaturated phospholipids, which was recapitulated by transduction of control neurons with 92 G_4_C_2_ repeats and prevented by treatment with a C9 ASO, demonstrating a disease-specific lipid signature. Promoting lipid desaturation through neuronal desaturase overexpression prolonged survival of C9 flies and rescued glutamate-induced excitotoxicity in C9 patient iPSC-derived spinal neurons. C9 flies also displayed a dramatic defect in cold stress recovery, a measure of impaired membrane fluidity, which was strongly reversed by neuronal desaturase overexpression.

Growing evidence links dysregulated lipid homeostasis to neurodegenerative diseases, including FTD/ALS. Several studies have shown altered levels of lipid species in FTD/ALS patient post-mortem tissue (*44*), cerebrospinal fluid (*45, 46*), and blood (*47–49*), as well as ALS rodent models (*50*). PUFAs have been specifically linked to ALS pathogenesis, with multiple epidemiological studies suggesting a protective role for dietary PUFAs in decreasing risk of developing ALS (*27, 28, 51*). A recent study of plasma fatty acids from 449 ALS patients revealed that higher levels of plasma α-linolenic acid at baseline are associated with prolonged survival and slower functional decline, while increased plasma linoleic acid and eicosapentaenoic acid (C20:5) were associated with a reduced risk of death during follow-up (*26*). Linoleic acid and α-linolenic acid are essential PUFAs that must be obtained from the diet, and that serve as precursors for the highly unsaturated species arachidonic acid (C20:4), eicosapentaenoic acid (C20:5), and docosahexaenoic acid (C22:6) (*52*). Here, we report that phospholipids containing these highly polyunsaturated fatty acids are specifically altered in C9 iPSC-neurons and FTLD post-mortem frontal cortex.

Our data fit well with the epidemiological evidence of PUFA levels and intake being protective in ALS, but crucially suggest that delivery of PUFAs to neurons is a key determinant of their protective function. We were able to study C9 lipid dysregulation specifically in neurons, by using an inducible neuronal driver in *Drosophila* and employing pure neuronal iPSC cultures for lipidomic analyses. Using this approach, we observed a strikingly enhanced benefit of neuronal overexpression of desaturases in flies versus feeding PUFAs in the diet. To reach the brain, PUFAs need to pass the gut barrier, as well as the blood brain barrier (BBB), and therefore the absolute quantities that reach neurons from the diet are unclear. A metabolic labelling study recently suggested that dietary sources account for 60-70% of the PUFAs in the mouse brain (*53*). However the efficiency of BBB transport varies for each fatty acid species (*54*). This delivery issue may explain the differences in survival benefits observed between genetic overexpression of desaturases versus pharmacological supplementation of their fatty acid products, although we cannot rule out that dietary linoleic acid and α-linolenic acid may partially mediate their survival benefits through systemic actions.

Aging is a major risk factor for both ALS and FTD (*55, 56*). Interestingly, overexpressing either *FASN1* or *Desat1* in neurons also significantly increased wildtype lifespan, suggesting that this pathway may also be beneficial to aging neurons and warrants further investigation in this context. Furthermore, we observed lipid-related transcriptional dysregulation and decreased PUFA-containing phospholipids not only in our C9 models but also in non-C9 FTD/ALS postmortem material. While our data show that C9 repeats are sufficient to drive lipid saturation changes, there must be other parallel pathways that induce these changes in sporadic forms of the disease, and aging-related changes are an obvious candidate. Taken together, impaired lipid metabolism is a common dysregulated pathway in FTD/ALS and it will now be important to investigate the different drivers of lipid-related changes.

The saturation of a lipid is determined by the number and position of double bonds in its esterified fatty acyl chains. Saturation, along with fatty acyl chain length and head group composition, influences membrane physicochemical properties and physiological functions (*57, 58*). The role of unsaturated lipids in modulating membrane fluidity has been well described (*38, 59–63*). Our study explored membrane fluidity in a physiological paradigm by testing the ability of flies to recover from cold stress. Flies neuronally expressing (G_4_C_2_)_36_ were sensitive to cold stress, which was ameliorated by overexpressing either *Desat1* or *FAT-2* with the same neuronal driver, suggesting that these desaturase enzymes are fluidizing neuronal membranes. Future studies are now warranted to assess the role of neuronal membrane fluidity in neurodegeneration. In addition to altering membrane dynamics, other mechanisms may also be involved. For example, PUFAs can be de-esterified from membrane phospholipids and converted to bioactive lipid mediators (*64–67*). Elevated levels of arachidonic acid-derived eicosanoids have previously been reported in ALS motor neurons, while inhibiting their production through 5-LOX inhibition has been shown to rescue toxicity in the developing eye in a C9 *Drosophila* model (*33*), thus highlighting another PUFA-related pathway that may contribute to disease.

While we focus here on neuronal lipids, future work may benefit from expanding these studies to glial and co-culture paradigms to unpick the interplay between different cell types. Indeed, recent work demonstrated that reactive astrocytes secrete saturated fatty acids, which promote motor neuron degeneration in ALS models (*68–71*), while astrocyte-specific knockout of *ELOVL1*, an enzyme responsible for producing long-chain saturated lipids, reduced astrocyte-mediated neuronal toxicity *in vitro* and *in vivo* (*68*). These data are in line with our findings as they converge on the hypothesis that PUFAs are protective to neurons while saturated fatty acids are harmful, which further highlights an important role for lipid desaturation in FTD/ALS pathogenesis. Overall, the results presented here identify dysregulated lipid metabolism as a direct contributor to neuronal toxicity in C9FTD/ALS and suggest that modulating neuronal lipid saturation is a promising approach for ameliorating C9-associated neurodegeneration.

## Methods

### *Drosophila* maintenance

*Drosophila* stocks were maintained on SYA food (15 g/L agar, 50 g/L sugar, 100 g/L autolysed yeast, 30 ml/L nipagin [10% in ethanol], and 3 ml/L propionic acid) at 25°C in a 12-hr light/dark cycle with 60% constant humidity. For RU486 induced experiments, food was supplemented with 200 µM RU486 (mifepristone). The elavGS stock was derived from the original elavGS 301.2 line (*72*) and generously provided by Hervé Tricoire (CNRS, France) (*73*). UAS-FASN1 and UAS-FASN2 lines were a gift from Jacques Montagne (Université Paris-Sud) (*74*). The w1118 line (BDSC:3605), was obtained from the Bloomington *Drosophila* Stock Centre. The UAS-Desat1 (DGRC:118679) and UAS-FAT-2 (DGRC:118682) lines were obtained from the KYOTO *Drosophila* Stock Centre (*35*). The UAS-(G_4_C_2_)_36_ stock has been previously described (*9*). All stocks were backcrossed to the w1118 strain for six generations before using for experiments.

### Fatty acid supplementation to *Drosophila* food

Fatty acids were added to SYA food, along with 200 µM RU486 (Sigma), while it was still liquid but had cooled to 50°C. The food was mixed thoroughly with an electric handheld blender, before dispensing into individual vials. Fatty acids used were palmitic acid (W283215, Merck), stearic acid (10002390, Fisher Scientific), oleic acid (W281506, Merck), linoleic acid (436305, Merck) and α-linolenic acid (L2376, Merck).

### *Drosophila* behavioural and lifespan assays

#### Lifespan assays

Flies were reared at a standard density in 200 mL bottles on SYA medium at 25°C. The parental generation was allowed to lay for 24-hr on grape-agar plates supplemented with yeast paste. Eggs were washed briefly in 1X PBS (pH 7.4) before being dispensed into bottles using a pipette at a standard density (20 µL of eggs in PBS, approximately 300 eggs). Two days post-eclosion flies were allocated to experimental vials at a density of 15 flies per vial (total of 150 flies per condition) containing SYA medium with or without 200 µM RU486. Deaths were scored and flies tipped onto fresh food three times a week. All lifespans were performed at 25°C on mated females. Data are presented as survival curves, and comparison between groups was performed using a log-rank test.

#### Assessment of Drosophila feeding

2-day-old mated female flies were transferred to SYA food containing 200 μM RU486 or ethanol vehicle control with PUFAs at a density of 5 per vial on the evening before the assay. Vials were coded and placed in a randomized order in rows on viewing racks at 25°C overnight. A feeding event was scored when a fly had its proboscis extended and touching the food surface while performing a bobbing motion. At the end of the assay, the vial labels were decoded, and the feeding data expressed as a proportion by experimental group (sum of scored feeding events divided by total number of feeding opportunities, where total number of feeding opportunities =number of flies in vial×number of vials in the group×number of observations) (*75*). For statistical analyses, comparisons between experimental groups were made on the totals of feeding events by all flies within a vial, to avoid pseudoreplication.

#### Cold stress recovery assay

*Drosophila* were exposed to 4°C for 18 hours using a modified protocol previously described (*41*). Recovery from cold-induced paralysis was scored after one hour at room temperature. The number of flies exhibiting a full recovery (walking), partial recovery (partial paralysis) or death were quantified and expressed as a percentage of total. The results were analyzed by χ2 test.

### *Drosophila* RNA sequencing sample and library preparation

Adult female flies were induced on SYA medium containing 200 µM RU486 or ethanol vehicle control and subsequently snap frozen. Total RNA was isolated from 15 heads per replicate using Trizol, and the experiment was performed in quadruplicate. RNA sequencing was performed with an Illumina NextSeq2000, using16 million paired-end reads/sample and 100 bp read length. Raw sequence reads were aligned to the Dm6 reference genome. DESeq2 (default parameters) was used to perform differential expression analysis. The “runTest” function from topGO package (v2.53.0) (*76*) was used to perform GO enrichment analysis on differentially expressed genes (|log2FoldChange| > 0.58). The “weight01” algorithm and “fisher” statistic were used when running topGO. The “GenTable” function was used to generate a table with the top biological process GO terms. Plots with top GO terms were plotted using ggplot2 (v3.4.2). We generated a heatmap for top GO terms showing the percentage of significantly differentially expressed genes among all genes of a GO term expressed in a dataset using the pheatmap function from pheatmap package (v1.0.12, https://CRAN.R-project.org/package=pheatmap).

### qRT-PCR

Total RNA was extracted from 15 heads per replicate, as above. Approximately 1 μg of RNA per sample (10.6 µL) was incubated with 2 µL TURBO DNase (Thermo Fisher Scientific) and 1.4 µL TURBO DNase buffer (Thermo Fisher Scientific) at 37°C for 15 minutes. Following this, the reaction was inhibited with addition of 2 µL EDTA to a final concentration of 3.4 mM, followed by incubation at 75°C for 5 minutes. 2 µL 0.5 µg/ µL oligo dT and 2 µL dNTP mix (10 mM stock made from individual 100 mM dNTP stocks, Invitrogen) were added to each sample followed by a 5-minute incubation at 65°C, after which samples were placed on ice. To each reaction, the following was added: 8 µL 5X first-strand buffer, 8 µL 25 mM MgCl_2_, 4 µL 0.1 M DTT, 2 µL RNaseOut RNase inhibitor (40 units/µL), 1 µL SuperScript II reverse transcriptase (Invitrogen). Samples were incubated at 42°C for 50 minutes, then heat inactivated at 70°C for 15 minutes. Quantitative PCR was performed using the QuantStudio 6 Flex Real-Time PCR System (Applied Biosystems) using SYBR® Green master mix (Applied Biosystems). Relative mRNA levels were calculated relative to alphaTub84B expression by the comparative Ct method. Primer sequences used are described in Table 2.

**Table 1.** Genotypes of stocks used throughout.

| Stock | Genotype |
| --- | --- |
| w<sup>1118</sup> | w<sup>[1118]</sup> |
| V-w<sup>+</sup> | w<sup>v<sup>1</sup></sup> |
| elavGS | w<sup>[1118]</sup>; P{elavGSGAL4} |
| UAS-(G<sub>4</sub>C<sub>2</sub>)<sub>36</sub> | w<sup>[1118]</sup>; P{UAS-GGGGCC.36}attP40 |
| UAS-FASN1 | w<sup>[1118]</sup>; P{UAS-FASN1.G} |
| UAS-FASN2 | w<sup>[1118]</sup>; P{UAS-FASN2.G} |
| UAS-Desat1 | w<sup>[1118]</sup>; P{w<sup>[+mC]</sup>=UAS-Desat1.S}16 |
| UAS-FAT-2 | w<sup>[1118]</sup>; P{w<sup>[+mC]</sup>=UAS-Cefat-2.S}3 |

**Table 2.**
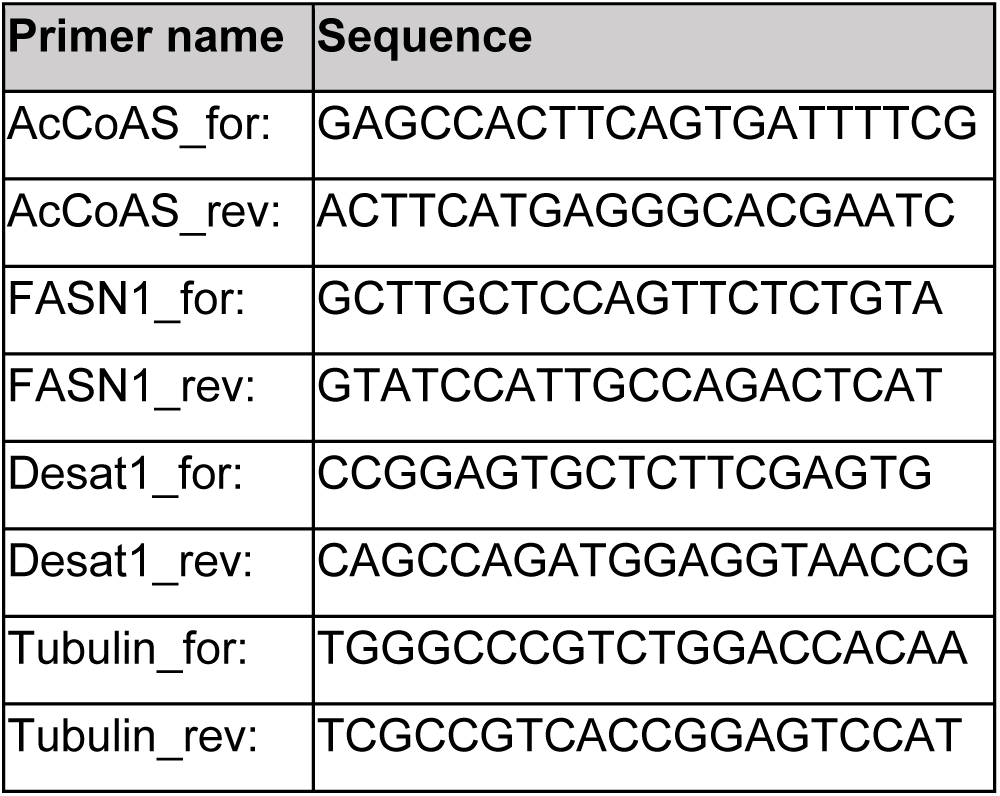
Primer sequences.

### DPR MSD immunoassays

#### Drosophila head protein preparation

Flies were set up as described for lifespan assays at a standard density and allowed to mate for 48 hours before being split onto control food or food supplemented with RU486 for 7 days. Following this, flies were snap frozen in liquid nitrogen. Heads were removed, and 10 heads per sample were homogenised in 100 μL 2% SDS buffer (Cat No. 428018, Merck) containing 1X RIPA buffer (Cat No. R0278, Sigma-Aldrich) and complete mini EDTA-free protease inhibitor cocktail (Cat No. 11836170001, Roche) at room temperature for about 30 sec until the heads were no longer intact. Samples were then heated at 95°C for 10 min. After centrifugation at 14,000 rpm for 20 min at room temperature, the supernatants were collected in the new tubes. The protein concentration was determined using Pierce BCA Protein Assay Kit (Cat No. 23325, ThermoFisher) according to the manufacturer’s manual.

#### i^3^Neuron protein preparation

i^3^Neuron replicates for DPR MSDs were collected alongside those used for lipidomic analyses from the same neuronal inductions. One well of a 6-well plate was used per replicate for MSD. At DIV21, neurons were lifted with PBS, centrifuged and pelleted at 1500xg for 5-10 min, snap frozen on dry ice, and stored at -80°C until use. For protein preparation, cell pellets were resuspended in 200 μL 2% SDS buffer (Cat No. BP2436-200, ThermoFisher) containing 1X RIPA buffer (Cat No. R0278, Sigma-Aldrich) and cOmplete mini EDTA-free protease inhibitor cocktail (Cat No. 11836170001, Roche) and sonicated 2x10 sec at 30 amp at 4°C. Sonicated samples were centrifuged at 17,000 g for 20 min at 16°C, after which supernatants were collected and used in MSD assays.

#### Running MSD assays

Samples were diluted to the same concentration with homogenisation buffer and 25 μL (fly samples) or 90 µL (cell samples) were loaded in duplicate in the 96-well Meso Scale Discovery (MSD) immunoassay plate. The singleplex MSD immunoassay was performed to measure poly(GA) or poly(GP) expression levels as described previously (*77*). The following antibodies were used: anti-poly(GP) (GP658, custom-made from Eurogentec) and anti-poly(GA) (clone 5E9, MABN889, Merck Millipore) as capture antibodies, and biotinylated anti-poly(GP) (GP658*) and biotinylated anti-poly(GA) (GA5F2*, kindly provided by Prof. Edbauer, Ludwig-Maximilians-Universität, München, biotinylated in house) as detector antibodies. Plates were read with the MSD reading buffer (R92TC; MSD) using the MSD Sector Imager 2400. A four-parameter logistic regression curve was fit to the values obtained from a standard curve using GraphPad Prism, and concentrations were interpolated. Signals correspond to the intensity of emitted light upon electrochemical stimulation of the assay plate. Before analysis, the average reading from a calibrator containing no peptide was subtracted from each reading.

### i^3^Neuron differentiation

C9 patient and isogenic control iPSC lines were kind gifts of the Chandran laboratory at University of Edinburgh (*43*) and C9 repeat knock-in lines on the KOLF2.1J background were a gift from the Skarnes laboratory at Jackson Labs as part of the iPSC Neurodegenerative Disease Initiative (iNDI) (*78*) (line details in Table 3). From these, we generated i^3^-compatible iPSC lines via piggyBac-integration of a BFP-containing, doxycycline-inducible *Neurogenin2* (Ngn2) minigene. After integration, iPSCs were subsequently doubly selected with puromycin and fluorescence-activated cell sorting, resulting in a pure population of stably expressing iPSCs which were expanded for use in these assays. These i^3^iPSCs were then used for rapid differentiation into cortical neurons (i^3^Neurons) using a previously described method (*30, 31*). Briefly, i^3^iPSCs were grown to 70-80% confluency, washed with PBS, lifted with Accutase (Gibco), and plated at 375,000 cells/well of a 6-well plate onto Geltrex-coated plates (DIV0). Cells were maintained from DIV0-3 in an induction medium consisting of DMEM-F12 (Gibco), 1x N2 (Thermo Fisher Scientific), 1x Glutamax (Gibco), 1x HEPES (Gibco), 1x Non-essential amino acids (Gibco), doxycycline (2 µg/mL) and 10 µM Y-27632 (DIV0 only; Tocris) which was exchanged daily. On DIV3, cells were dissociated with accutase and replated onto poly-L-ornithine (Merck) or polyethyleneimine (Sigma) and laminin (Sigma) coated 6-well plates at 600,000 cells/well in neuronal maintenance media consisting of Neurobasal (Gibco), supplemented with 1x B27 (Gibco), 10 ng/mL BDNF (PeproTech), 10 ng/mL NT-3 (PeproTech) and 1 µg/mL laminin. From DIV3 to DIV21, cells were maintained in neuronal maintenance media, with ⅓ media changes once weekly. Lentiviral transduction to overexpress (G_4_C_2_)_92_ or (G_4_C_2_)_2_ was done 1 hour after DIV3 replating. Likewise, antisense oligonucleotide (ASO) treatments to target the *C9orf72* sense strand or a non-targeting control were also begun on DIV3 and supplemented in media changes thereafter. In brief, 1 hour after replating, ASOs were transiently transfected using Lipofectamine Stem (Invitrogen STEM00015) at 5 μM final concentration according to manufacturer’s protocol. 1 day after ASO treatment, a full media change was done to remove remaining Lipofectamine Stem and replaced with neuronal maintenance media containing 5 μM ASO which was then further re-supplemented in weekly media changes at 5 μM. ASOs were published in (*7*) and have fully modified phosphorothioate backbones. The sequences are as follows, with the five 2’ O-Methyl RNA basepairs on either end (italicized):

C9 sense-targeting: *UACAG*GCTGCGGTTG*UUUCC*

Non-targeting: *CCUUC*CCTGAAGGTT*CCUCC*

**Table 3.** i^3^iPSC line information.

| iPSC line name | Common name | Sex (M/F) | Clinical diagnoses | Number of G<sub>4</sub>C<sub>2</sub> repeats | Source of iPSCs |
| --- | --- | --- | --- | --- | --- |
| BS6 | C9orf72 line 1 | F | ALS/FTD | ~750 | Chandran lab, Univ. of Edinburgh |
| BS6-2H9<br>(Isogenic control of BS6) | Control line 1 | F | N/A | Repeats removed by CRISPR-Cas9 | Chandran lab, Univ. of Edinburgh |
| WTc11 | Control line 2 | M | N/A | N/A | Commercial |
| KOLF2.1J-D08 | C9orf72 line 2 | M | N/A | KOLF2.1J with ~224 repeats knocked-in | iNDI (78, 79) |
| KOLF2.1J-F05 | C9orf72 line 3 | M | N/A | KOLF2.1J with ~202 repeats knocked-in | iNDI (78, 79) |
| KOLF2.1J | Control line 3 | M | N/A | Parental line | iNDI (78, 79) |

### (G_4_C_2_)_92_ or (G_4_C_2_)_2_ lentiviral construct subcloning

pCDH-EF1-MCS-IRES-copGFP lentiviral plasmid (System Biosciences) was used as the backbone to create (G_4_C_2_)_92_ and (G_4_C_2_)_2_ lentiviral constructs. Subcloning to insert the repeats was undertaken in a two-step process. First, we synthesized a DNA fragment consisting of a custom multiple cloning site (MCS) sandwiched in between 300 bp each of repeat-adjacent upstream and downstream sequence from *C9orf72* intron 1 and then inserted it into the internal MCS of pCDH-EF1-MCS-IRES-copGFP with InFusion cloning (Takara Bio) in between XbaI and NotI restriction sites. This interim construct, termed “pCDH-EF1-C9up-MCS-C9down-IRES-copGFP” was verified with diagnostic restriction digests and Sanger sequencing across the insert. Then, to create the (G_4_C_2_)_92_ construct, a 92-repeat sequence was isolated from a previously verified in-house construct with NheI and NotI restriction digests and subcloned into the MCS of pCDH-EF1-C9up-MCS-C9down-IRES-copGFP with overnight ligation at 4°C (T4 ligase, NEB). To maintain repeat stability, bacterial clones were grown at room temperature, in half the standard ampicillin concentration (0.5 mg/mL), and in low-salt LB broth (Sigma). A repeat-stable clone was selected and subsequently maxi-prepped (Qiagen) for use in lentiviral production. Thus, the final construct consisted of 92 repeats immediately surrounded on either side by 300 bp of endogenous *C9orf72* intronic sequence to facilitate RAN translation and upstream of an IRES-copGFP sequence for fluorescent visualization of transduction efficiency. To create the (G_4_C_2_)_2_ control lentiviral constructs, two complementary short oligos were synthesized containing 2 G_4_C_2_ repeats and NheI and NotI restriction site overhangs. Oligos were resuspended in annealing buffer (NEB buffer 2.1), heated to 95°C, and allowed to cool slowly to room temperature to anneal. Annealed oligos were used directly in ligation reactions into pCDH-EF1-C9up-MCS-C9down-IRES-copGFP with the same protocol as used for the 92-repeat construct.

### Lentiviral production

HEK293T cells were grown at 37°C and 5% CO_2_ in T175 flasks. At ∼70% confluency, cells were transfected with either (G_4_C_2_)_92_ and (G_4_C_2_)_2_ lentiviral transfer plasmids along with PAX (Addgene #12260) and VSV-G (Addgene #12259) lentiviral packaging plasmids with Lipofectamine 3000 Transfection Reagent (Invitrogen) according to manufacturer protocol. 48 hours later, media was collected and centrifuged at 500 g for 10 min at 4°C to remove cell debris, after which Lenti-X Concentrator (Takara Bio) was added at a 1:3 ratio. After a minimum incubation of 24h at 4C, concentrator/media mix was centrifuged at 1500xg at 4°C for 45 min and resulting concentrated lentiviral pellet was resuspended in sterile PBS, aliquoted, and stored at -80°C until use.

### Targeted lipidomics of i^3^Neurons and post-mortem brain samples

#### Sample collection

At DIV21, i^3^Neurons were pelleted and stored at −80°C until analysis. Brains were donated to the Queen Square Brain Bank (QSBB) (UCL Queen Square Institute of Neurology) with full, informed consent. Clinical and demographic data for all brains used in this study were stored electronically in compliance with the 1998 data protection act and are summarised in Table 4. Ethical approval for the study was obtained from the NHS research ethics committee (NEC) and in accordance with the human tissue authority’s (HTA’s) code of practice and standards under licence number 12198. All cases underwent a pathological diagnosis for FTLD according to current consensus criteria (*80, 81*). The cohort included pathologically diagnosed cases of FTLD without *C9orf72* expansion (n=33), FTLD with *C9orf72* expansion (n=15) and neurologically normal controls (n=13). Frontal cortex grey matter and cerebellum was dissected from each brain and stored at − 80°C until analysis.

**Table 4.** Clinical and demographic data for post-mortem brain samples.

| Group | Sample size | Age at onset (Avg. +/- SD) | Age at death (Avg. +/- SD) | % Male | Pathological Diagnosis | Mutations |
| --- | --- | --- | --- | --- | --- | --- |
| Control | n=13 | N/A | 74 ± 6.1 | 53.8% | N/A | N/A |
| FTLD-non-C9 | n=32 | 60 ± 8.5 | 70 ± 7.6 | 59.3% | FTLD | GRN (n=8)<br>none (n=24) |
| FTLD-C9 | n=15 | 57.7 ± 6.2 | 65.3 ± 7.0 | 60.0% | FTLD | C9orf72 het (n=13)<br>C9orf72 homo (n=1)<br>C9orf72 het + GRN (n=1) |

#### Targeted lipidomic measurements

Comprehensive targeted lipidomics was accomplished using a flow-injection assay based on lipid class separation by differential mobility spectroscopy and selective multiple reaction monitoring (MRM) per lipid species (Lipidyzer platform; SCIEX, Framingham, USA). A very detailed description of lipid extraction, software and the quantitative nature of the approach can be found elsewhere (*82–84*). In short, after the addition of >60 deuterated internal standards (IS), lipids were extracted using methyl tert-butyl ether. Organic extracts were combined, dried under a gentle stream of nitrogen, and reconstituted in running buffer. Lipids were then analyzed using flow-injection in MRM mode employing a Shimadzu Nexera series HPLC and a Sciex QTrap 5500 mass spectrometer. For the internal calibration, deuterated IS lipids for each lipid class were used within the lipidomics workflow manager. Each lipid species was corrected by the closest deuterated IS within its lipid class and afterwards the obtained area ratio was multiplied by the concentration of the IS.

### Analyses of targeted lipidomic datasets

#### Filtering and normalizations

Raw amounts of individual lipid species were obtained from the Lipidyzer platform as above and subsequently filtered and normalized. Datasets were first filtered for low-abundance and undetected lipid species. To pass filtering, a lipid species must be detected in at least 80% of all samples in the analysis as well as 60% of samples in any given group and must also be at least 2-fold above the average of the blanks. After filtering, missing sample values were imputed as the median of other samples in their group; this step was found to be necessary for subsequent normalizations, as missing values greatly skewed the proportional datasets. Next, filtered, and imputed datasets were normalized either to total lipids (for analysis of class-level lipid alterations, as in **Sup Fig. 6, 8**) or by lipid class individually (for analysis of saturation shifts and indices, as in **Fig. 2B-C and Fig. 3B-C**). Thus, these processing steps result in proportional lipidomic measurements, relative to either the total lipidome or total amount of lipid within each class, respectively.

#### Fold-changes and unsaturation indices in i^3^Neurons

For these analyses, “biological replicates” were considered as different i^3^Neuron lines and “technical replicates’’ were individual cell pellets from those lines that were grown, collected, and analyzed separately. Technical replicates were obtained across a minimum of 2-3 neuronal inductions per line or condition. To assess changes in saturation and calculate unsaturation indices, we utilized lipid class-normalized data from these technical replicates. To attain fold-changes for i^3^Neurons, as shown in the heatmap in **Fig. 2B** and **Sup Fig. 3**, each technical replicate was filtered and normalized individually. Fold-changes were then calculated for each lipid species as the average amount across technical replicates in an induction over the average in its internal control condition (e.g. replicates of control lines with LV-(G_4_C_2_)_92_ were compared to those with LV-(G_4_C_2_)_2_ in the same induction). **Fig. 2C** displays average fold-changes across inductions for each line/condition over control. The heatmap in **Sup Fig. 3** displays fold-changes for each induction individually, in order to demonstrate the reproducibility of the loss of highly unsaturated phospholipids across experiments. To calculate the unsaturation index, a composite score was calculated for each technical replicate individually, using the ratio of the sum of phospholipid species with 4-6 double bonds in their most highly unsaturated fatty acyl chain over the sum of species with 0-3 double bonds. Technical replicates were then combined across biological replicates (i.e. across all three C9 lines) and unpaired Student’s t-tests were used to compare the experimental and control conditions, as in **Fig. 2C**. Normality tests were performed for each condition with D’Agostino & Pearson’s test which determined each distribution to be normally distributed (α=0.05).

#### Fold-changes and unsaturation indices in post-mortem brain samples

Fold-changes in FTLD versus control samples, as used in the heatmap in **Fig. 3B**, were calculated for each lipid species separately as the average of the FTLD condition over the average of the control condition, using lipid class-normalized data. Unsaturation indices were calculated as was done for i^3^Neurons, considering each brain sample as a replicate, and unpaired Student’s t-tests were used to compare the FTLD and control conditions.

#### Volcano plots for i^3^Neurons and post-mortem brain samples

To generate volcano plots for i^3^Neurons, technical replicates were combined within each biological replicate separately (e.g. C9 line 1) and fold-changes and p-values were obtained using unpaired Student’s t-tests on lipid class-normalized datasets. Moreover, fold-changes and p-values for the post-mortem lipidomic datasets between FTLD and control conditions were computed.

### Glutamate-induced excitotoxicity assays in iPSC-derived spinal neurons

#### Subcloning for BFP, FAT-1 and FAT-2 overexpression constructs

mTagBFP (Addgene #89685), FAT-1 (Genscript), and FAT-2 (Genscript) cDNAs were amplified with PCR and subcloned into the pHR-hSyn-EGFP vector (Addgene #114215) along with a T2A-NLS-mApple minigene for fluorescent visualization. In brief, EGFP was removed with BamHI and NotI (NEB) and BFP/FAT-1/FAT-2 and T2A-NLS-mApple fragments were inserted with InFusion cloning (Takara Bio), as per manufacturer’s protocol. Resulting plasmids were verified with diagnostic restriction digest and Sanger sequencing prior to being maxi-prepped (Qiagen) for subsequent use in excitotoxicity assays.

#### Excitotoxicity assays

Non-neurological control and *C9orf72* iPSCs were obtained from the Answer ALS repository at Cedars Sinai (see **Table 5** for demographics) and maintained in mTeSR Plus medium at 37°C with 5% CO_2_. iPSC derived spinal neurons (SNs) were differentiated according to a modified diMNs protocol (*85–88*) and maintained at 37°C with 5% CO_2_. iPSCs and iPS-SNs were routinely tested negative for mycoplasma. On day 12 of differentiation, iPS-SNs were dissociated with Trypsin. 5 x 10^6^ iPS-SNs were nucleofected with 4 µg plasmid DNA in suspension as previously described (*86, 87*). Following nucleofection, 100 µL of cell suspension was plated in each well (total of 6 wells per cuvette) of a glass bottom or plastic 24 well plate for PI and Alamar blue toxicity and viability experiments respectively. Media was exchanged daily for a total of 20 days to facilitate the removal of iPS-SNs that failed to recover post-nucleofection. On the day of the experiment (day 32 of differentiation), iPS-SN media was replaced with artificial CSF (ACSF) solution containing 10 µM glutamate. For those iPS-SNs undergoing Alamar Blue viability assays (plastic dishes), Alamar blue reagent was additionally added to each well according to manufacturer protocol at this time. Following incubation, iPS-SNs for PI cell death assays were incubated with PI and NucBlue live ready probes for 30 min and subjected to confocal imaging as previously described (*87*). The number of PI spots and nuclei were automatically counted in FIJI. Alamar Blue cell viability plates were processed according to manufacturer protocol. As a positive control, 10% Triton X-100 was added to respective wells 1-hr prior to processing.

**Table 5.** Demographic Information for iPSN lines.

| iPSC line name | Age at collection | Sex (M/F) | Clinical diagnosis | Source |
| --- | --- | --- | --- | --- |
| CS0201 | 56 | F | Non-neurologic control | Cedars-Sinai |
| CS0002 | 51 | M | Non-neurologic control | Cedars-Sinai |
| CS0206 | 72 | F | Non-neurologic control | Cedars-Sinai |
| CS9XH7 | 53 | M | Non-neurologic control | Cedars-Sinai |
| CS8PAA | 58 | F | Non-neurologic control | Cedars-Sinai |
| CS1ATZ | 60 | M | Non-neurologic control | Cedars-Sinai |
| CS0NKC | 52 | F | C9orf72 | Cedars-Sinai |
| CS0LPK | 67 | M | C9orf72 | Cedars-Sinai |
| CS0BUU | 63 | F | C9orf72 | Cedars-Sinai |
| CS7VCZ | 64 | M | C9orf72 | Cedars-Sinai |
| CS6ZLD | Unknown | F | C9orf72 | Cedars-Sinai |
| CS8KT3 | 60 | M | C9orf72 | Cedars-Sinai |

### Statistical Analysis

The statistical test used for each experiment is indicated in the figure legends. Log-rank tests for fly survival were performed in Microsoft Excel (template described in (*89*)). ANOVA or Student’s t-test analyses were performed in GraphPad Prism v10.0.2. For all statistical tests, p<0.05 was considered significant.

## Acknowledgements

This work was supported by funding from Alzheimer’s Research UK (AMI and LP), the European Research Council (ERC) under the European Union’s Horizon 2020 research and innovation programme (648716—C9ND) (AMI), the UK Dementia Research Institute, which receives its funding from UK DRI Ltd, funded by the UK Medical Research Council, Alzheimer’s Society, and Alzheimer’s Research UK (AMI), EMBO (AJC), the Live Like Lou Foundation (AJC), the Chan Zuckerberg Initiative (AMI, RvdK, MG), NIH NINDS R01 NS132836 (ANC) and NIH NINDS/NIA R00 NS123242 (ANC). The Queen Square Brain Bank for Neurological Disorders at University College London Queen Square Institute of Neurology receives support from the Reta Lila Weston Institute for Neurological Studies.

