## Supplemental Figures for "Neuronal polyunsaturated fatty acids are protective in FTD/ALS"

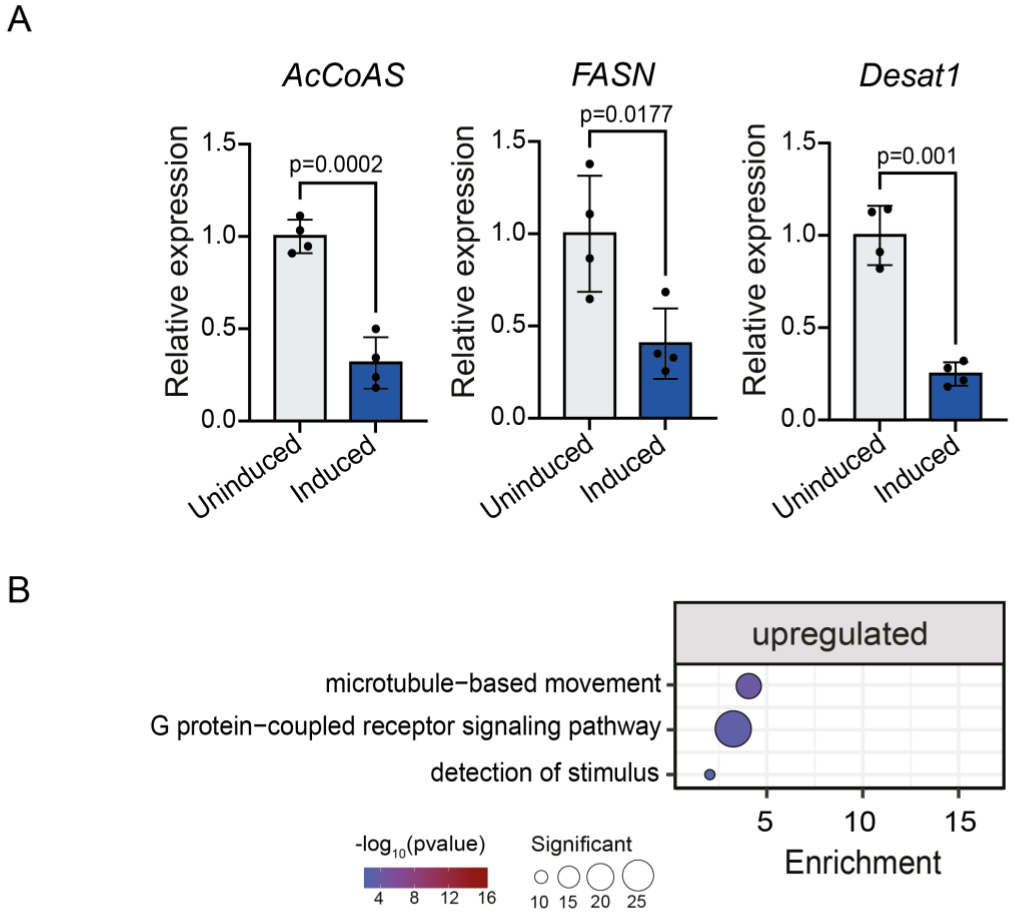


**Supplementary Figure 1. RT-qPCR validation of differentially expressed genes in C9 flies versus controls and gene ontology (GO) enrichment analyses of upregulated genes.** (A) Confirmation of C9 *Drosophila* RNA-seq result by RT-qPCR, showing significant downregulation of *AcCoAS, FASN1* and *Desat1* in C9 *Drosophila* heads versus controls, normalised to tubulin. (B) GO enrichment of upregulated genes in C9 flies versus controls.

A

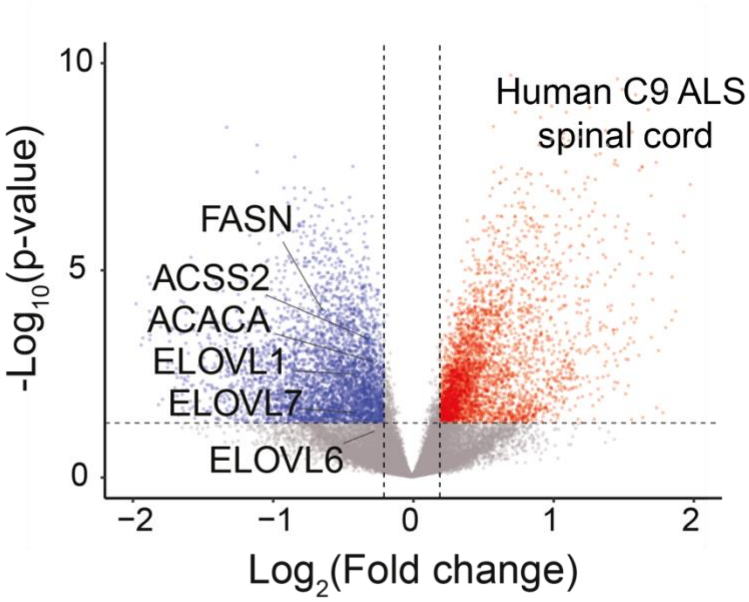


**Supplementary Figure 2. Fatty acid synthesis and desaturation pathway genes are downregulated in C9 ALS post-mortem cervical spinal cord.** (A) Volcano plot of RNA-seq data from cervical spinal cord of 28 patients with C9 ALS and 36 non-neurological disease controls from the New York Genome Center ALS Consortium (*29*) highlighting canonical fatty acid synthesis and desaturation pathway genes as downregulated.

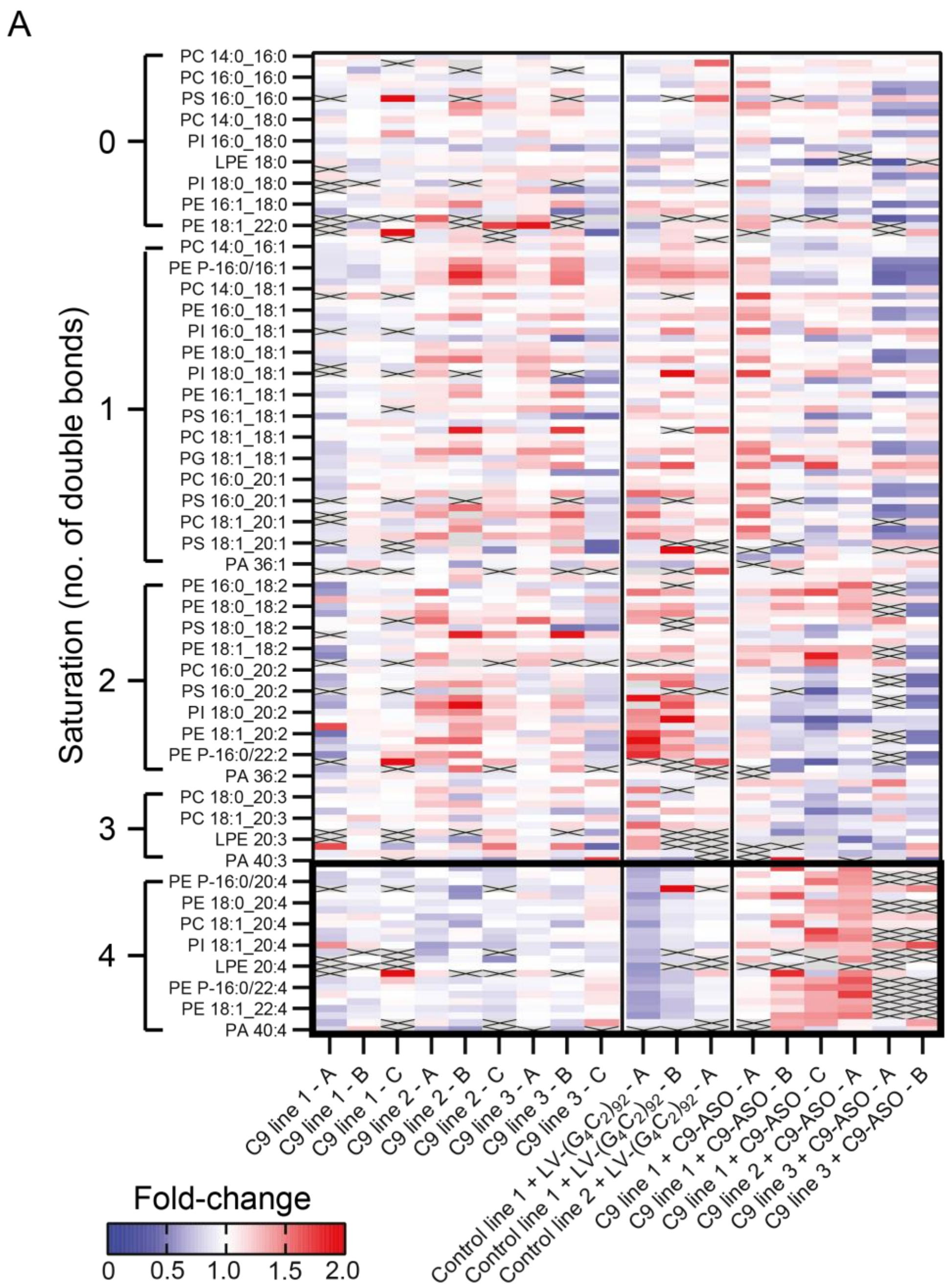


**Supplementary Figure 3. Phospholipid levels in i[3]Neurons, displayed as separate neuronal inductions.** (A) Heatmap displaying all detected phospholipids as fold-change over control in each neuronal induction separately. Lipids are normalized by lipid class. Grey boxes indicate phospholipid species that are outside the fold-change range, while X's represent species which were not detected. Loss of highly unsaturated species is consistently observed across neuronal inductions in C9 lines and control lines expressing 92-repeats, while phenotype is prevented by C9-ASO treatment.

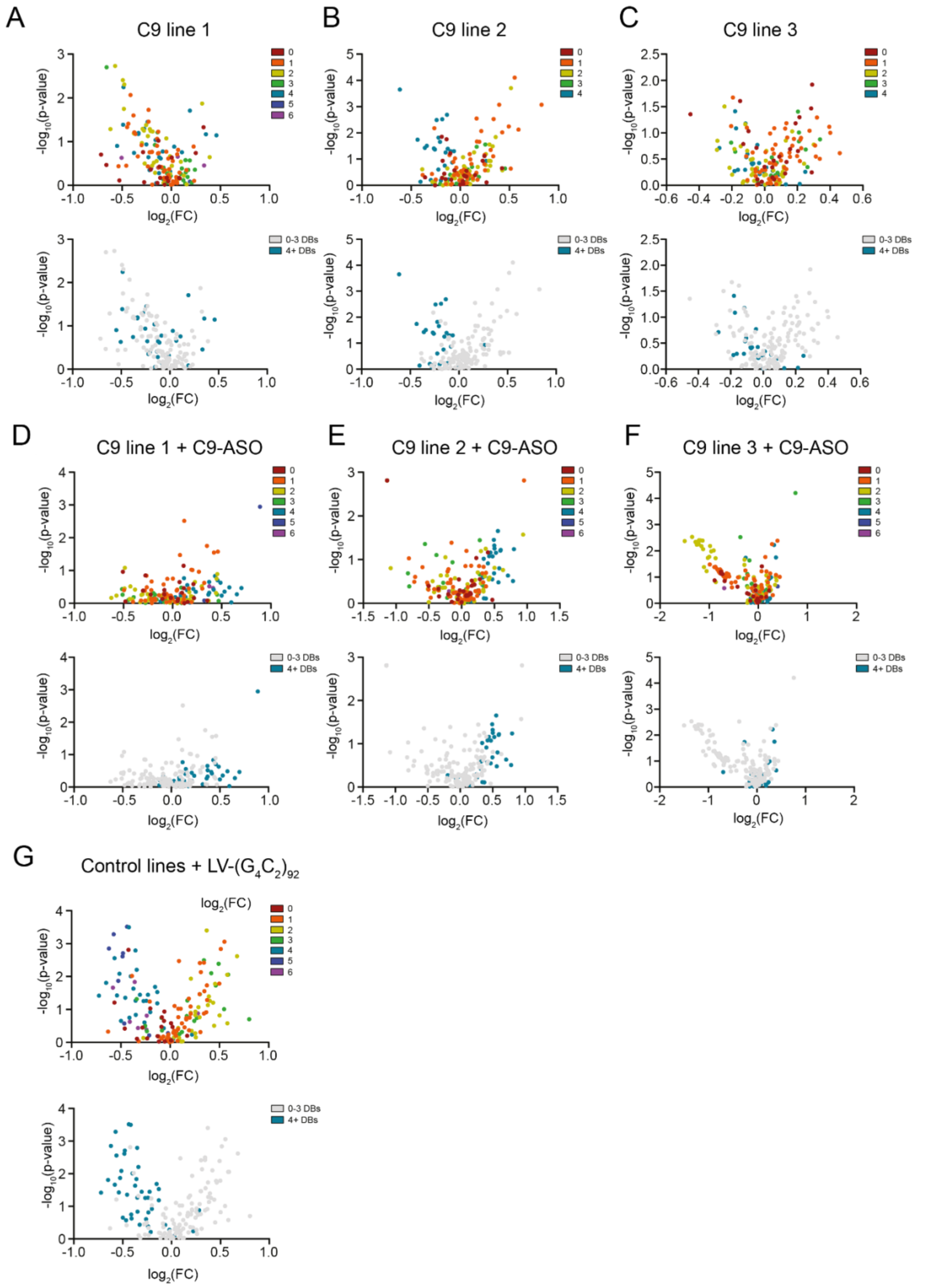


**Supplementary Figure 4. Volcano plots of phospholipids in C9 i[3]Neuron lines compared to isogenic controls and lines treated with sense repeat (C9) targeted antisense oligonucleotides (ASOs) or LV-($G_4C_2$)$_{92}$.** (A-C) Volcano plots of all detected phospholipid species in each C9 line compared to its individual isogenic control line, displaying downregulation of highly unsaturated species (≥4 double bonds). Values represent $log_2$(fold change over control) and significance (Student’s t-test) across all replicates/inductions within the labeled group. In top plots, color corresponds to the number of double bonds in the species’ most unsaturated fatty acyl chain. (D-F) Volcano plots displaying all phospholipid species in each C9 line treated with a C9-ASO compared to a NT-ASO control. (G) Volcano plots of phospholipid species in control lines treated with LV-$(G_4C_2)_{92}$ compared to a LV-$(G_4C_2)_2$ control.

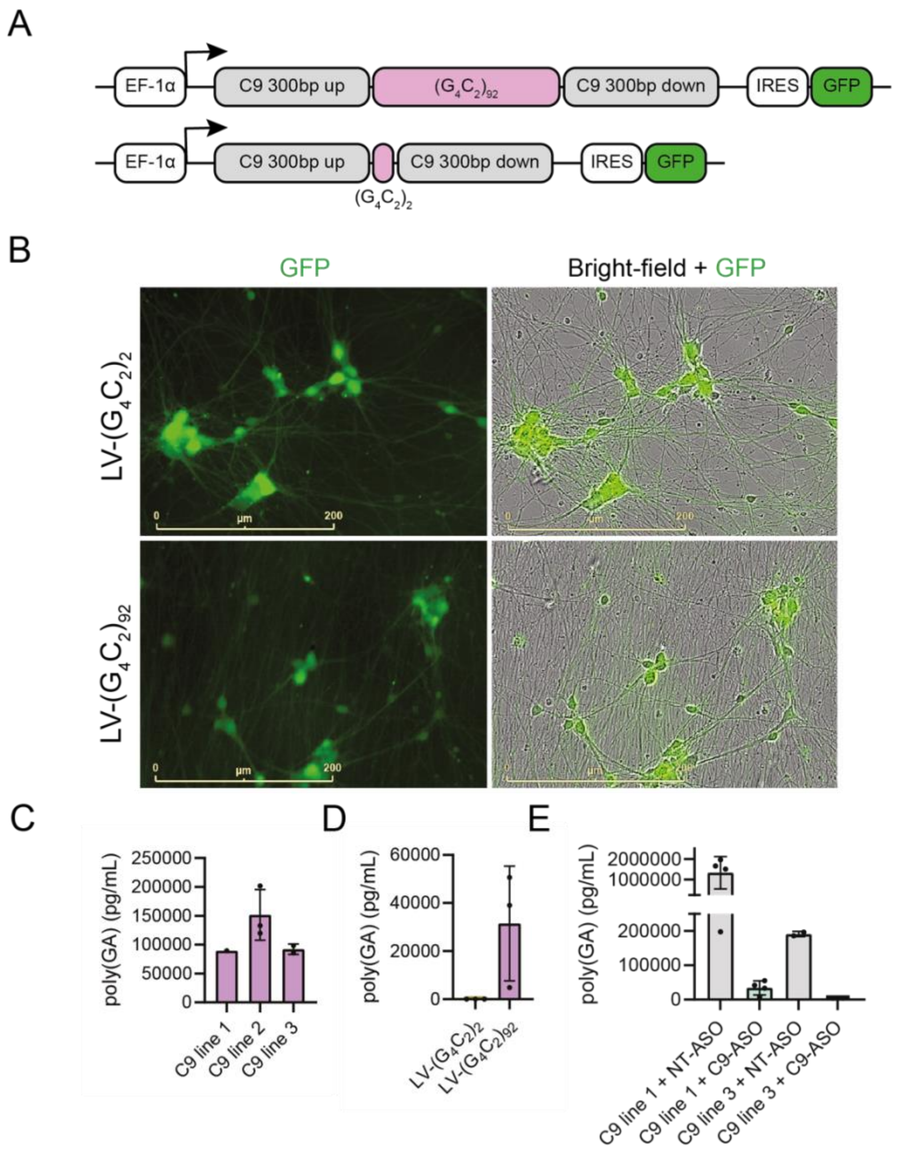


**Supplementary Figure 5. poly(GA) levels in i[3]Neuron lines treated with ($G_4C_2$) lentiviruses or sense repeat-targeted antisense oligonucleotides (ASOs).** (A) Lentiviral 92 (LV-$(G_4C_2)_{92}$) and 2 (LV-$(G_4C2)_2$) repeat constructs have 300 bp of endogenous repeat-flanking sequence to facilitate RAN translation and include an IRES-GFP for live-cell visualization. (B) LV-$(G_4C_2)_{92}$ and LV-$(G_4C_2)_2$ lentiviruses were titrated via GFP signal (live imaged) to high transduction efficiencies for lipidomic experiments. (C-E) poly(GA) immunoassay in (C) C9 lines, (D) control lines treated with LV-$(G_4C_2)_{92}$ or LV-$(G_4C_2)_2$, and (E) C9 lines treated with sense repeat-targeted (C9) ASOs or non-targeted (NT) control ASOs from (*7*). All measurements were taken on DIV21 from the same neuronal inductions used for lipidomic analyses.

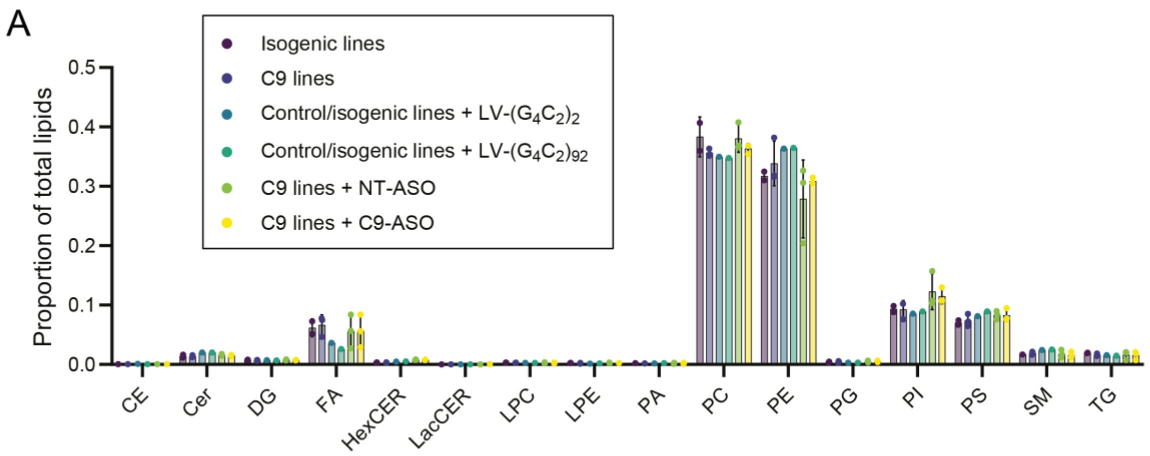


**Supplementary Figure 6. Lipid class distributions in i[3]Neurons.** (A) Lipid classes displayed as proportion of total lipidome in i[3]Neurons. Dots represent biological replicates (i.e. independent i[3]Neuron lines). CE = cholesterol ester; Cer = ceramide; DG = diacylglyceride; FA = fatty acid; HexCER = hexosylceramide; LacCER = lactosylceramides; LPC = lysophosphatidylcholine; LPE = lysophosphatidylethanolamine; PA = phosphatidic acid; PC = phosphatidylcholine; PE = phosphatidylethanolamine; PG = phosphatidylglycerol; PI = phosphatidylinositol; PS; phosphatidylserine; SM = sphingomyelin; TG = triacylglyceride.

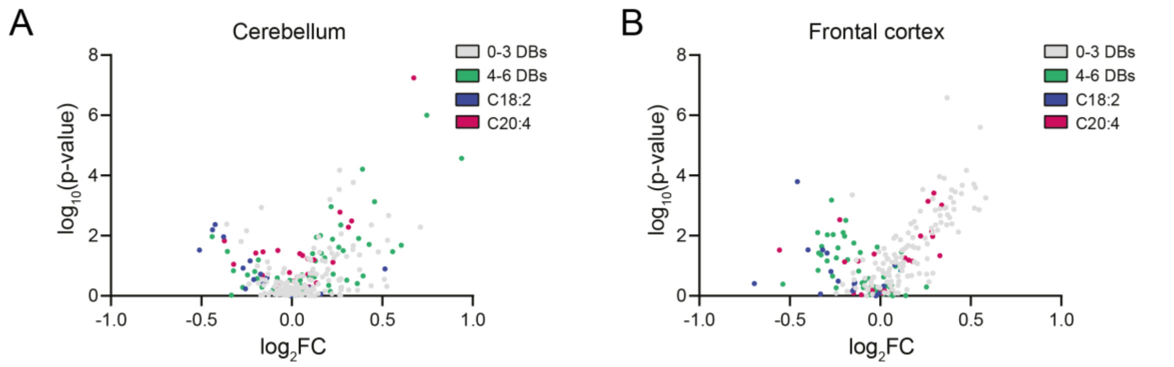


**Supplementary Figure 7. Volcano plots of phospholipid species in FTLD versus control post-mortem tissues.** (A-B) Volcano plot of all detected phospholipid species in FTLD compared to non-neurological control (A) cerebellum and (B) frontal cortex, displaying downregulation of highly unsaturated species (≥4 double bonds) in the frontal cortex. Values represent $log_2$(fold change over control) and significance (Student's t-test).

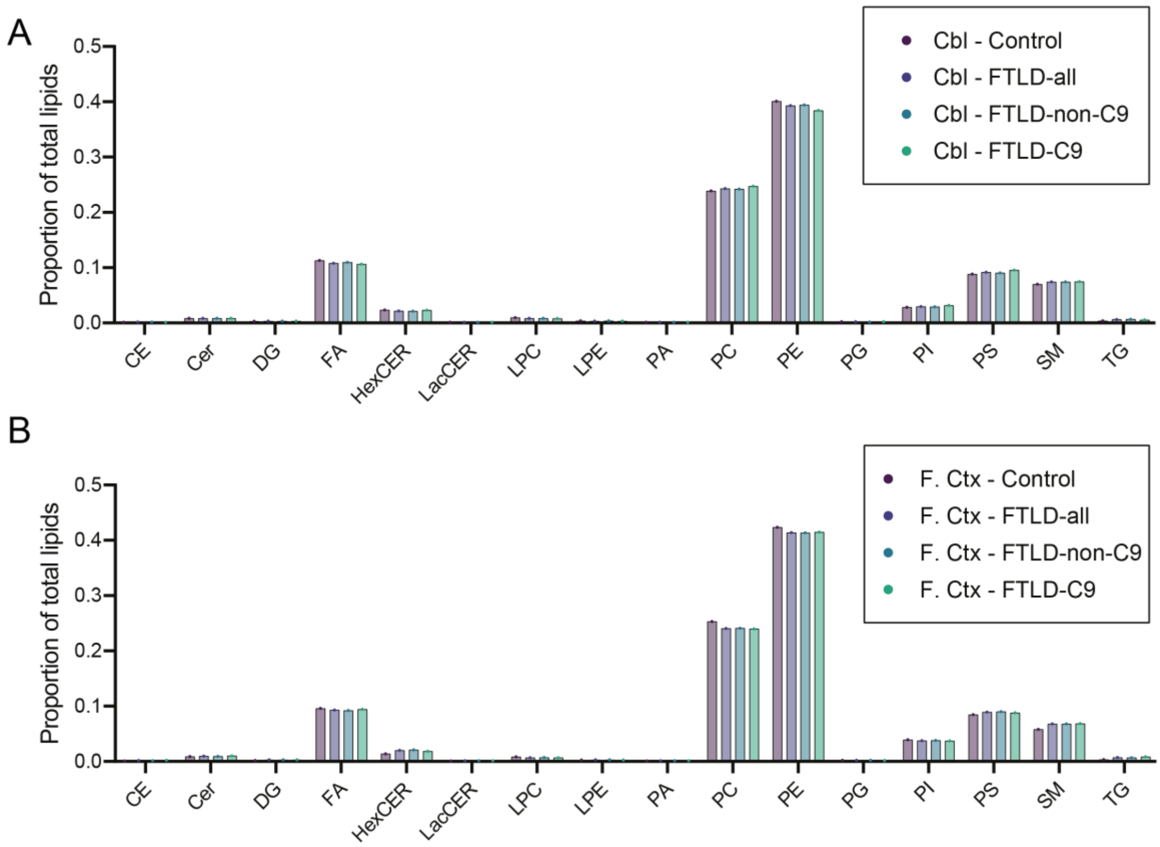


**Supplementary Figure 8. Lipid class distributions in FTLD post-mortem tissues.** (A) Lipid classes displayed as proportion of total lipidome in control and FTLD post-mortem cerebellum and frontal cortex. Bars represent average across all samples (n=46-48 FTLD, n=13 control). CE = cholesterol ester; Cer = ceramide; DG = diacylglyceride; FA = fatty acid; HexCER = hexosylceramide; LacCER = lactosylceramides; LPC = lysophosphatidylcholine; LPE = lysophosphatidylethanolamine; PA = phosphatidic acid; PC = phosphatidylcholine; PE = phosphatidylethanolamine; PG = phosphatidylglycerol; PI = phosphatidylinositol; PS; phosphatidylserine; SM = sphingomyelin; TG = triacylglyceride.

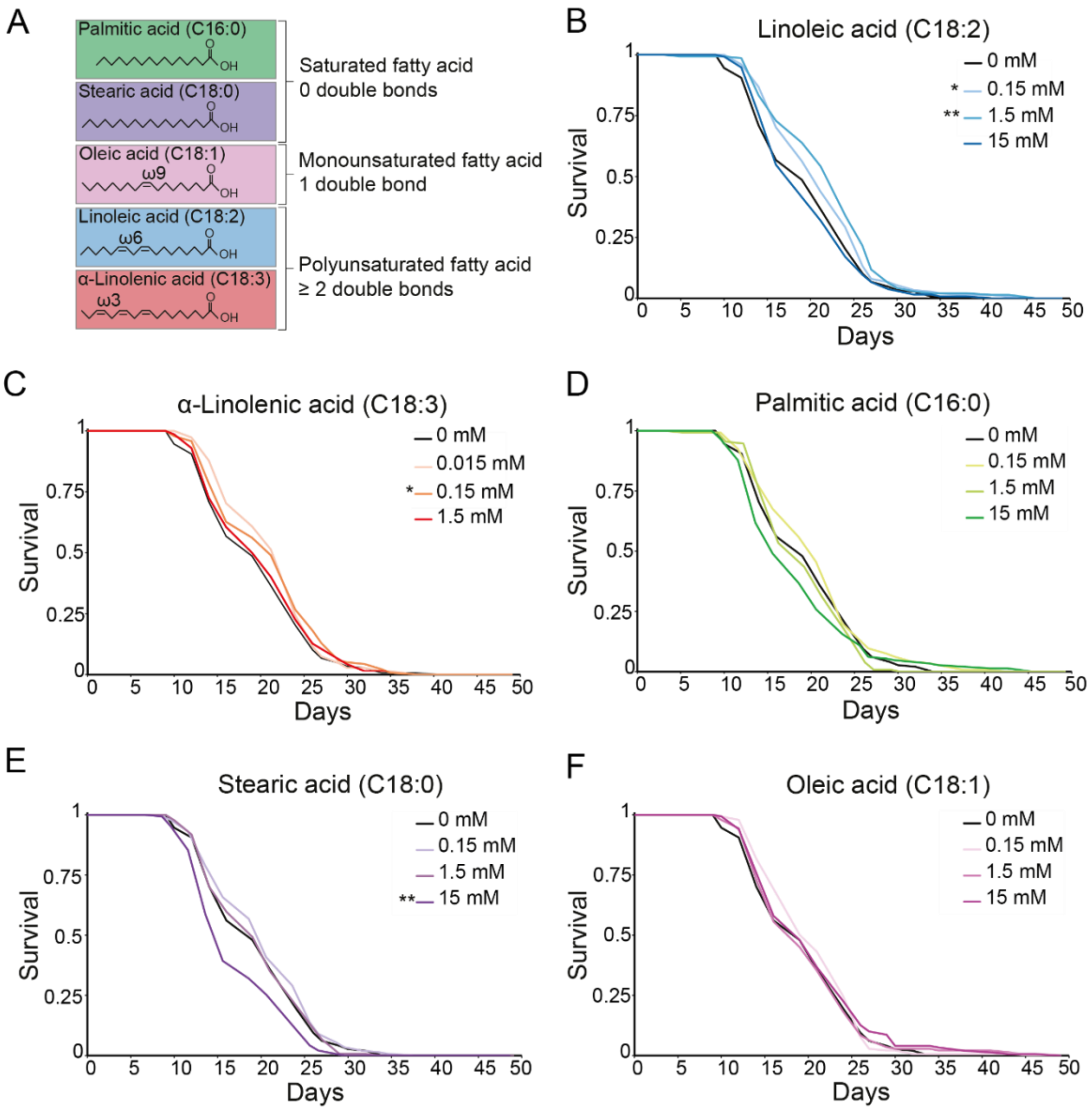


**Supplementary Figure 9. Lifespans of C9 flies fed saturated, monounsaturated, and polyunsaturated long chain fatty acids.** (A) Structure and saturation of selected fatty acids. (B) Palmitic acid had no significant effect on survival of C9 flies at any of the concentrations tested (0.15 mM, p=0.052, 1.5 mM p=0.182, 15 mM p=0.473). (C) Stearic acid had no significant effect at 0.15 mM (p=0.079) or 1.5 mM (p=0.992), but significantly decreased survival at 15 mM (p=0.008). (D) Oleic acid had no significant effect on survival at any of the concentrations tested (0.15 mM, p=0.285, 1.5 mM p=0.782, 15 mM p=0.186). (E) Linoleic acid significantly increased survival of C9 flies at 0.15 mM (p=0.036) and 1.5 mM (p=0.002) concentrations, but not 15 mM (p=0.663). (F) Supplementation of α-linolenic acid significantly increased survival of C9 flies at 0.15 mM (p=0.049), but not at 0.015 mM (p=0.055) or 1.5 mM (p=0.216). n=135-150 flies per condition. Log-rank test used for all comparisons. Genotype: UAS-($G_4C_2$)36, elavGS.

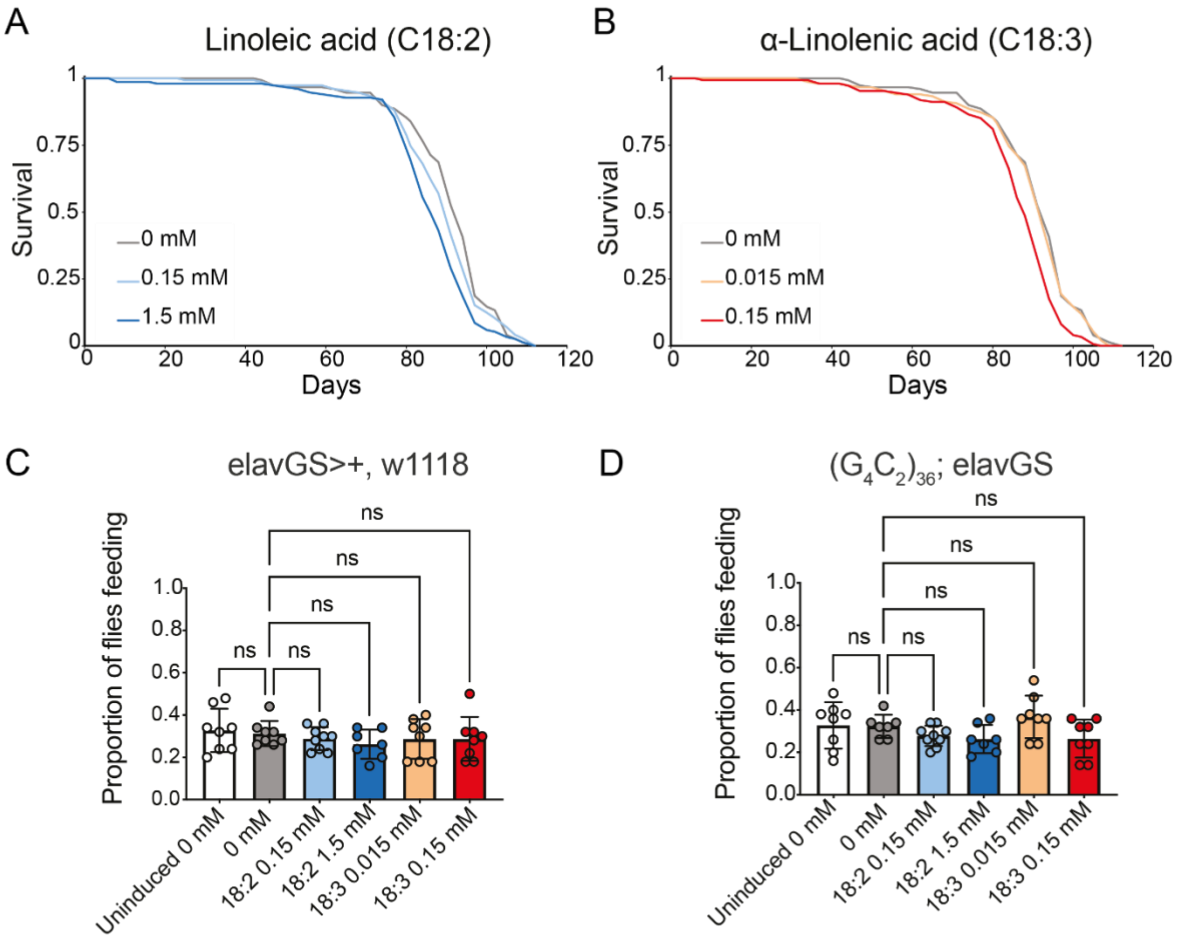


**Supplementary Figure 10. Linoleic and α-linolenic acid supplementation does not affect wildtype lifespan and does not alter proboscis extension response of wildtype or C9 flies.** (A) Linoleic acid supplementation had no effect on wildtype lifespan at 0.15 mM (p=0.162) and significantly shortened wildtype lifespan at the 1.5 mM ($p=4.352x10^{-5}$) concentration. (B) Supplementation with α-linolenic acid had no effect on wildtype lifespan at 0.015 mM (p=0.599), and significantly shortened wildtype lifespan at the 0.05 mM concentration ($p=6.22\times10^{-6}$), n=150 flies per condition. Log-rank test used for all comparisons. (C-D) Food supplementation with linoleic or α-linolenic acid does not alter proboscis extension response of wildtype or C9 flies. Flies were placed onto new food 24 hours before assay was performed, at a density of five flies per vial, n=7-9 biological replicates. All groups were induced with RU486 except for the uninduced conditions. Two-way ANOVA with Tukey's multiple comparison test was used to calculate statistical significance. Data presented as mean ± S.D. Genotypes: elavGS, UAS-$(G_4C_2)_{36}$, elavGS.

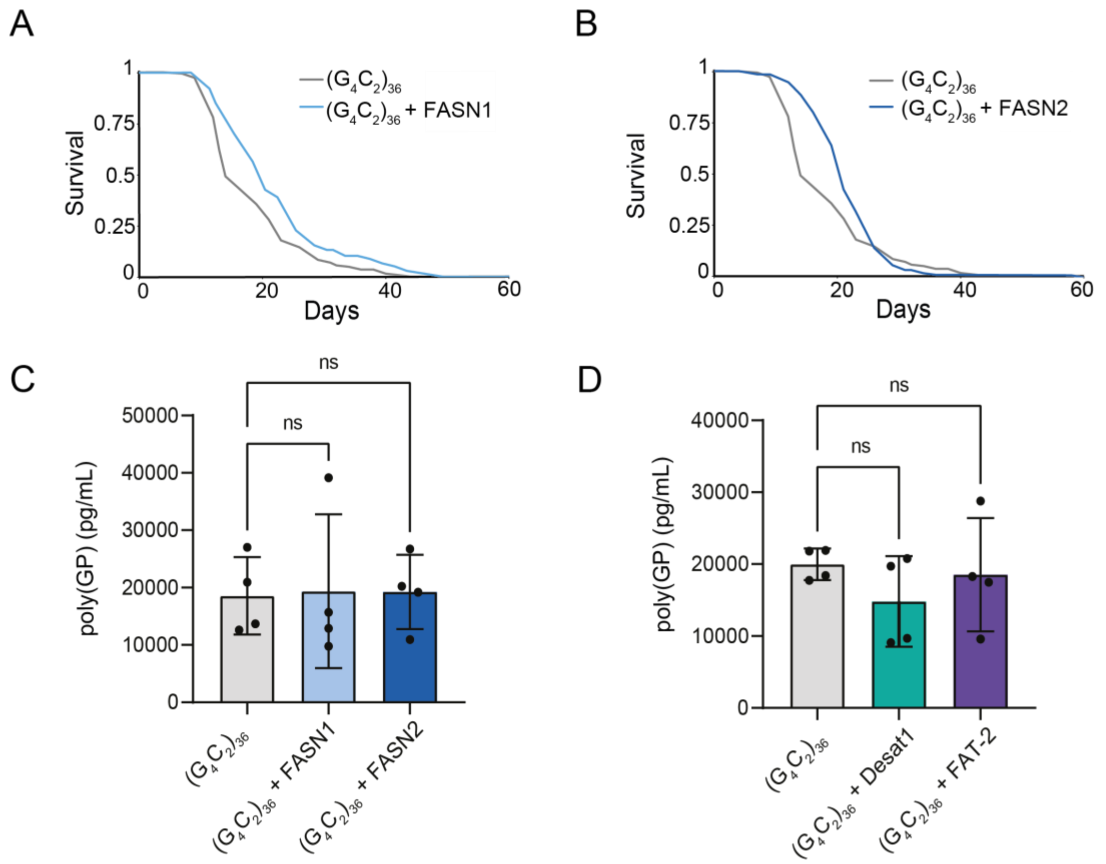


**Supplementary Figure 11. Overexpression of fatty acid synthases extends C9 survival, while fatty acid synthase and desaturase overexpression does not alter poly(GP) levels.** (A-B) Neuronal overexpression of *FASN1* or *FASN2* extended C9 fly survival (FASN1 $p=1.16\times10^{-5}$; FASN2 p=0.003), n=150 flies per condition, log-rank test used for each comparison. (C) Neuronal expression of *FASN1* or *FASN2* did not alter poly(GP) levels in $(G_4C_2)_{36}$ fly heads. (D) Neuronal expression of *Desat1*, or *FAT-2* did not alter poly(GP) levels in $(G_4C_2)_{36}$ fly heads. One-way ANOVA, followed by Tukey's post-hoc test. n=4 replicates per condition. Data presented as mean ± S.D. Genotypes: UAS-$(G_4C_2)_{36}$, elavGS; UAS-$(G_4C_2)_{36}$/UAS-FASN1, elavGS; UAS-$(G_4C_2)_{36}$/UAS-FASN2, elavGS; UAS-$(G_4C_2)_{36}$, elavGS/UAS-Desat1; UAS-$(G_4C_2)_{36}$, elavGS/UAS-FAT-2.

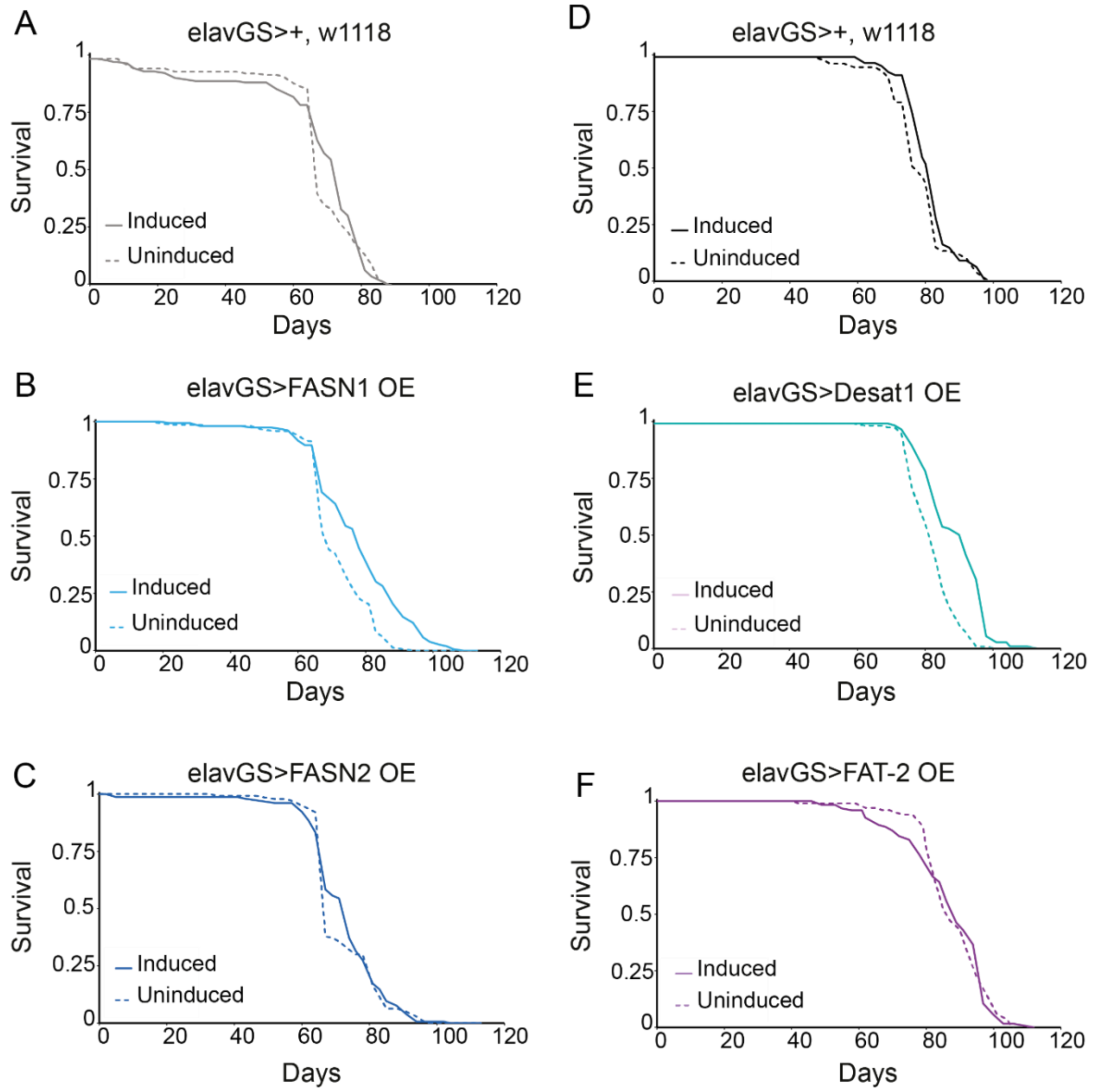


**Supplementary Figure 12. Lifespans of wildtype flies overexpressing fatty acid synthases or desaturases.** (A) Driver alone control lifespan run in parallel to B and C. There was no difference in lifespan between RU486 induced and uninduced flies (p=0.408). (B) *FASN1* overexpression in neurons of wildtype flies extended lifespan (p=$3.386\times10^{-8}$). (C) *FASN2* overexpression in neurons of wildtype flies had no effect on lifespan (p=0.866). (D) Driver alone control lifespan run in parallel to E and F. There is no difference in lifespan between RU486 induced and uninduced flies (p=0.197). (E) *Desat1* overexpression in neurons of wildtype flies extended lifespan (p=$1.567\times10^{-10}$). (F) Overexpression of *FAT-2* in neurons of wildtype flies had no effect on lifespan (p=0.589), n=150 flies per condition. Log-rank test used for all comparisons. Genotypes: elavGS; elavGS/UAS-FASN1; elavGS/UAS-FASN2; elavGS/UAS-Desat1; elavGS/UAS-FAT-2.
